# Lateralized Tenm4 expression biases visual performance and modulates axonal growth and pre-synapse size in retinal ganglion cells

**DOI:** 10.64898/2026.08.28.747855

**Authors:** Zexin Zhao, Andrea Szydlo-Shein, Léa Darnet, Katherine E. Trevers, Robert Hindges

## Abstract

The precise wiring of neuronal connections is a striking feature of the central nervous system and is critical for correct brain function. During vertebrate visual development, retinal ganglion cells (RGCs) establish synaptic connections with postsynaptic cells in the optic tectum, a process that is in part conferred by cell adhesion molecules. However, a complete understanding of the retinotectal wiring mechanism is lacking. Here, we show that the synaptic adhesion molecule Teneurin-4 (Tenm4) is expressed by RGCs and their postsynaptic tectal cell targets in a left-right side-specific manner in zebrafish. Loss of *tenm4* leads to defects in presynaptic structures in RGCs and further affects their axonal growth and arbor complexity. Functional assessment of RGC responses to visual stimuli showed a general increase in response amplitude in mutant animals. Visual behavioral analysis revealed a deficit with a strong left-right asymmetrical component in mutant larvae. Our findings demonstrate that *tenm4* plays an important role in sensorimotor circuit assembly, extending to neural activities and visual acuity.

## INTRODUCTION

The precise assembly of visual circuits from initial axonal extension to synaptic development allows the animal to interact with its environment and carry out specific visuomotor behaviors.^1–3^ This process is highly stereotypical during development across vertebrates.^4–6^ The primary output of the retina is established by retinal ganglion cells (RGCs), which send out axons towards the midline which then, depending on the degree of binocular overlap in the species, cross to the contralateral side or stay ipsilaterally and carry the sensory information to their principle midbrain target, the optic tectum (superior colliculus in mammals).^7–9^ Their elaborate axon arborization in this target area establishes synaptic connections with tectal cells (TCs).^10,11^ The establishment and refinement of synaptic connections are modulated by cell adhesion molecules (CAMs), which act as interaction partners or provide molecular cues for synaptic specificity crucial for function and execution of visual behaviours.^12–14^ However, our detailed understanding of the roles of CAMs in shaping these circuits and providing information for synapse formation remains incomplete.

Teneurins are evolutionarily conserved type II transmembrane glycoproteins that are widely expressed in multiple tissues, with the highest levels in brain.^15,16^ They are composed of a small N-terminal intracellular sequence, a single transmembrane domain, and a large extracellular region,^17,18^ through which it modulates synaptic specificity via homophilic and heterophilic molecular interactions.^19,20^ Four orthologs have been identified in vertebrates (*tenm1* to *tenm4*)^17,21^. Mutations in all human Teneurin genes have been associated with multiple neurodevelopmental and psychiatric disorders.^22–26^ Teneurin-4 contributes to embryonic patterning,^27^ oligodendrocyte differentiation and myelination,^28,29^ neurite outgrowth,^30^ and the formation of ipsilateral RGC projections.^31^ Since Tenm4, a prominent CAM in the visual system, is expressed along the visual pathway from early developmental stages,^32,33^ it is possible that it has a crucial role in RGC axon outgrowth and synapse formation, with potential consequences for visual function.

Here, using a site-specific knock-in approach, we find that Tenm4 is expressed across all retinal cells in zebrafish with a distinctive left-right asymmetry in the retinotectal circuit. Developmental analysis of single axons in a *tenm4* CRISPR knock-out (KO), including time-lapse imaging, reveals that Tenm4 plays a role in RGC axonal growth dynamics and loss of the protein alters the correct formation of presynaptic structures. In addition to the structural effects seen in mutant animals, we detect changes in the functional responses of RGC upon visual stimulation, as seen through glutamate release at individual synapses as well as in calcium responses on the population level. Behavioral assessments indicate a change in visual acuity in *tenm4* mutant animals, with specific lateralized defects. Our findings uncover an important function of Tenm4 during the initial circuit development with a specific role in presynaptic formation and link the morphological and synaptic phenotypes with sensorimotor responses.

## RESULTS

### Tenm4 is expressed across the body, displaying heterogeneity in the retinotectal circuit

Our prior work showed that Tenm4 is widely expressed in the developing zebrafish nervous system^32^. To better assess the individual cells expressing *tenm4* with an emphasis on sensorimotor circuits, we used a CRISPR knock-in strategy inserting the transcriptional activator Gal4 in-frame with the *tenm4* start codon at exon 2, followed by a P2A sequence to allow normal production of Tenm4 and maintain endogenous protein levels (*Tg(tenm4:Gal4)^kg1Tg^*, Figure 1A and S1B).^34^ For validation we used hybridization chain reaction - fluorescent RNA *in situ* (HCR-FISH). All HCR patterns and signals from the Gal4 reporter line were aligned to the brain coordinates of the mapzebrain atlas standart.^35,36^

**Figure 1.**
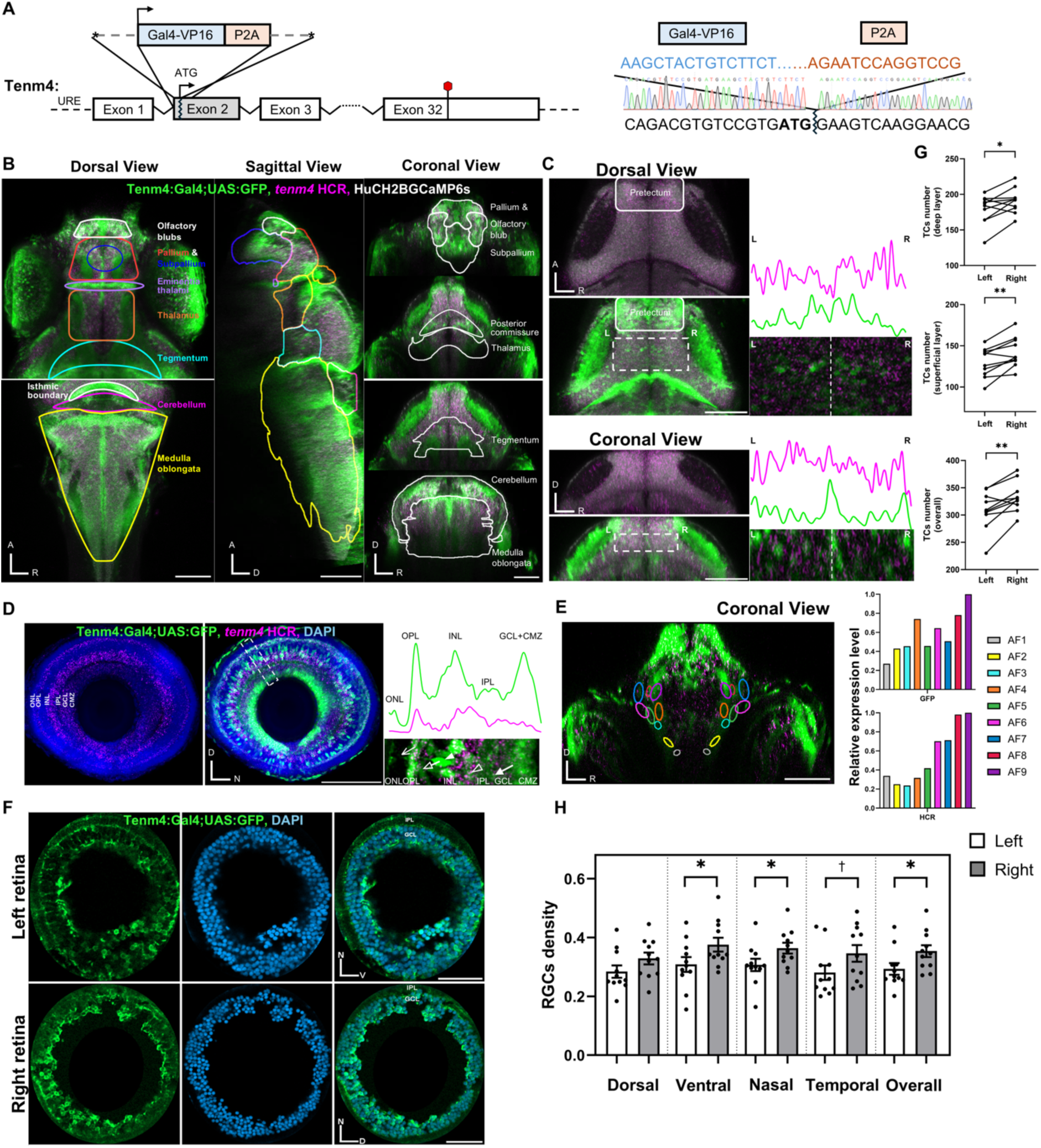
Tenm4 expression at RNA and cellular level in early development. **A)** Schematics of the CRISPR knock-in strategy of *Gal4* transcription factor followed by a P2A peptide sequence downstream of *tenm4* start codon at exon 2 (left) and sequence validation (right)URE: upstream regulatory element; *: 5’-biotin and phosphonothioate bond in the first 5 nucleotides; Red rhombus: termination site of translation. **(B to D)** Images from *tenm4* in situ experiments (magenta) and GFP-positive cells (green) from the *Tg(tenm4:Gal4*;*UAS:GFP)* line co-registered to the standard brain atlas at mapzebrain.org. The orthogonal views of Tenm4 expressed regions are outlined, including olfactory bulbs, pallium, subpallium, eminentia thalami, anterior and dorsal thalamus, posterior commissure, torus longitudinalis, medially dorsal tegmentum, isthmic boundary, cerebellum, medulla oblongata (B), as well as pretectum and optic tectum **(C)**, and retina **(D)**, with the fluorescent intensity profile (dashed white square) across retinal and tectal layers. White-filled arrow: RGCs; open arrowhead: ACs; white-filled arrowhead: BCs; non-filled arrow: HCs; open arrow: PRs. Scale bar: 100μm. IPL/INL/OPL/ONL: inner/outer plexiform/nuclear layer. GCL: ganglion cell layer. CMZ: ciliary marginal zone. **(E)** Coronal view of highlighted arborization fields (AFs) 1-9 in different colors and their relative expression level of *tenm4* GFP and HCR signals. Scale bar: 100μm. **(F)** Cryo-sectioned left and right retina from *Tg(tenm4:Gal4*;*UAS:GFP)* with DAPI staining (as reference). Scale bar: 40μm. **(G to H)** The number of tectal cells (TCs) are quantified in the left and right tectum at superficial and deep layer of the 20μm z-stack **(G)**. The percentage of RGCs density was bilaterally quantified in dorsal, ventral, nasal, temporal regions of left and right retina, averaged from one superficial and deep slice **(H)**. n=10 and p-values are from two-tailed paired t-test. D: dorsal; A: anterior; R: right; N: nasal; V: ventral.

Applying HCR in *Tg(tenm4:Gal4*;*UAS:GFP)* larvae (Figure 1B to 1D), we found that the mRNA signals mostly recapitulated reporter expression in the form of discrete puncta, both of which were enriched in the visual motor commands involved in the sensorimotor circuitry, including anterior and dorsal thalamus, posterior commissure, torus longitudinalis, the nucleus of medial longitudinal fasciculus, pretectum, optic tectum, cerebellum and nucleus isthmi. Most prominently, *tenm4* mRNA signals were highly enriched in layers containing neuronal soma. In the retina (Figure 1D), besides the previously described expression in the ganglion cell layer (GCL) and the inner nuclear layer (INL) region close to the inner plexiform layer (IPL) containing RGCs and amacrine cells, respectively,^32^ *tenm4* was also strongly detected across the entire INL while more sparsely in the outer nuclear layer (ONL), corresponding to the location of bipolar cells and horizontal cells as well as in the photoreceptors. Measuring the fluorescence profile of GFP positive arbors in the IPL showed that there is an even distribution between the ON and OFF sublaminae. At the distal retinal edge, we found abundant Tenm4-positive (Tenm4^+^) cells in the ciliary marginal zone (CMZ), a niche for stem and progenitor cells, suggesting a potential role in cell differentiation.^37,38^

There are 10 distinct retino-recipient areas in zebrafish, named arborization field (AF) 1-10.^39^ In the principal AF10 – the optic tectum – Tenm4^+^ tectal cells (TCs) showed a salt-and-pepper pattern within the tectal stratum, along with Tenm4^+^ RGC-axons innervating the neuropil, possibly indicating a regulatory role in neuropil stratification (Figure 1C). With regard to other AFs, Tenm4^+^ RGCs appeared to be preferentially restricted to the AF 6-9 (Figure 1E), the subregions of the pretectum.^40^ Notably, these regions are also innervated by Tenm4^+^ pretectal neurons, implicating Tenm4 in organizing circuits involved in looming-evoked and visuomotor behaviors.^40,41^ Taken together, *tenm4* is expressed all along the visual pathway, from the individual retinal neurons to various retino-recipient AFs and further downstream circuits, suggesting a possible role in instructing connectivity along the visual pathway to balance behavioral phenotypes.

When analyzing Tenm4^+^ (GFP-positive) cells across animals, we noticed an asymmetrical distribution between the left and the right side of the larvae. For in-depth analysis, we prepared cryo-sections from the *Tg(tenm4:Gal4*;*UAS:GFP)* retinae and used whole-mount tectal preparations to quantify the GFP-positive cells, with a general nuclear stain as reference for the retinal sections. Surprisingly, we found that Tenm4^+^ cell numbers were significantly biased towards the right animal side in both RGCs (i.e. left eye versus right eye) and TCs (i.e. left hemisphere vs. right hemisphere) (Figure 1C and 1F). RGCs are distributed as an average of 35.36% ± 2.03% on the right side versus 29.30% ± 1.99% on the left side (*p < 0.05*,* Figure 1G), when compared to total cell numbers. In the tectum (absolute cell numbers) we found an average of 333 ± 9 TC on the right side versus only 309 ± 11 TCs on the left side (*p < 0.01\*\**, Figure 1G and 1H). We next examined whether this laterality originates from specific retinal areas. To this end, we divided the retina into four quadrants: dorsal, ventral, nasal and temporal (Figure 1F). In line with previous *in situ* hybridization data,^32^ Tenm4^+^ neurons were uniformly distributed across all four quadrants within one eye (Figure 1H). However, when comparing the left and the right eye, we found a significant bias towards the right retina for the ventral and nasal parts and a strong trend for the temporal quadrant (difference: V: 6.73% ± 2.74%, *p < 0.05\**; N: 5.69% ± 2.43, *p < 0.05\**; T:6.54% ± 3.12%, *p = 0.0623*). In the optic tectum (Figure 1G), a similar bias towards the right side was observed in both the superficial (S) and deep (D) layers (difference: S: 13 ± 3, *p < 0.01\*\**; D: 12 ± 4, *p < 0.05\**, Figure 1H). These data suggest that *tenm4*^+^ RGCs and TCs are asymmetrically distributed between the left and right eyes/brain hemispheres but its relative proportions across cell types within a side stay the same.

### Tenm4 is required for morphological development of axons and presynaptic terminals in RGCs

To date, research on *tenm4* functions using genetic deletion models has primarily focused on overall circuit formation between neurons of different areas^42,43^ but has overlooked the effects in single neurons on the fine structure of synaptic organization. To investigate this, we created a *tenm4* knock-out mutant (*tenm4^kg^*^98^*^/kg^*^98^, hereinafter named as *tenm4^-/-^*) using CRISPR-Cas9 genome editing techniques. In *tenm4^-/-^* mutants, a 10bp nucleotide deletion within the sequence encoding the intracellular domain of *tenm4* results in a frameshift and, consequently, a premature stop codon, causing the loss of both the transmembrane and extracellular domains (Figure 2A and S1A). Quantitative PCR revealed that *tenm4* mRNA expression levels were diminished by ∼90% in *tenm4^-/-^* and ∼10% in *tenm4^+/-^* mutants, most likely due to nonsense-mediated RNA decay (Figure S1B).

**Figure 2.**
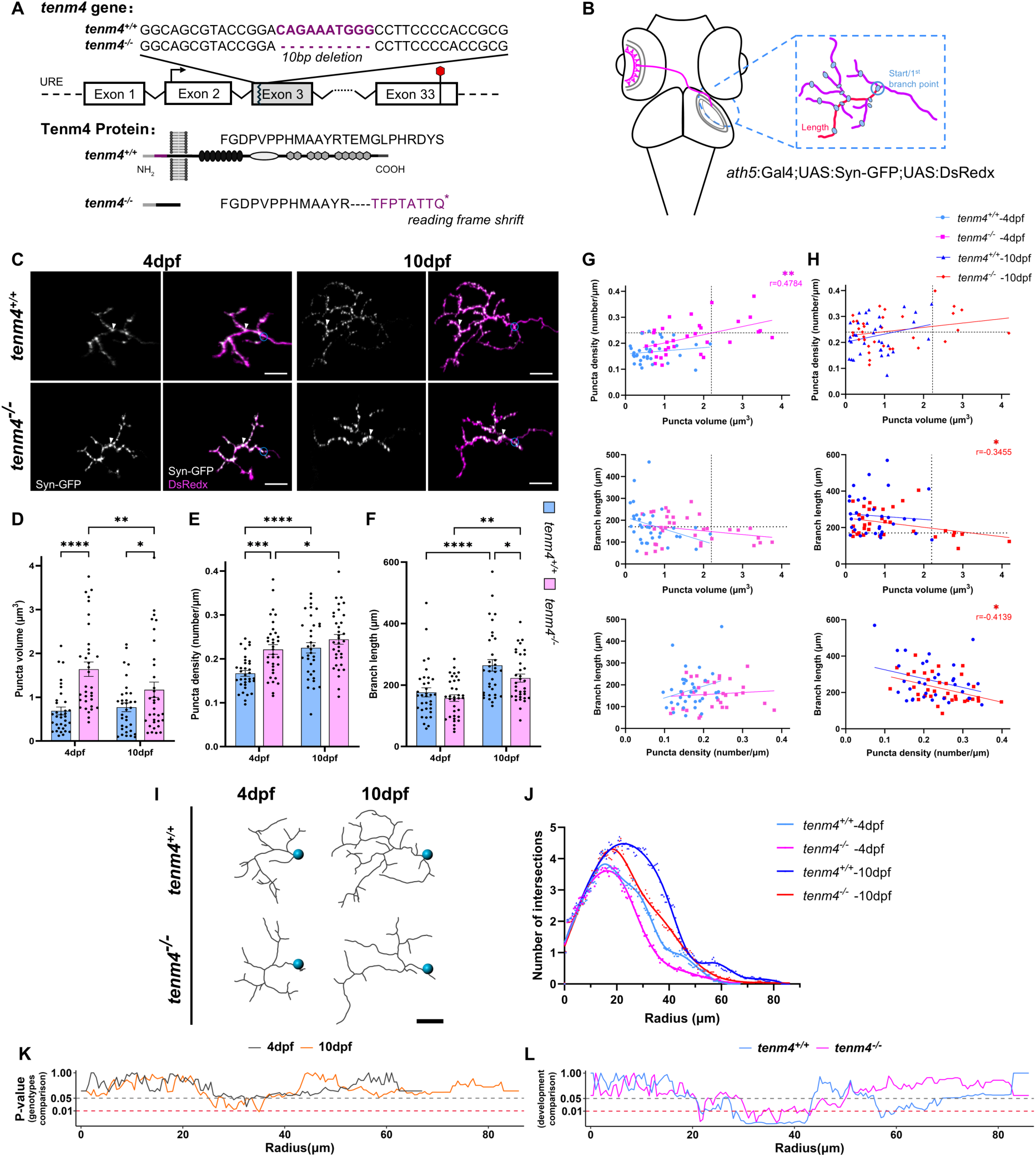
***Tenm4* mutants exhibit aberrant RGC presynaptic structures and axonal arborizations (A)** CRISPR/Cas9-mediated 10bp deletion in the *tenm4* gene leading to premature STOP and consequently loss of all extracellular domains in the Tenm4 protein (bottom). **(B)** Schematic showing sparse labelling strategy of RGC with presynaptic puncta and axonal arbor in the optic tectum, with the start point set at the 1^st^ branching point (indicated by blue circle) and branch length measurement. **(C)** Confocal images of the sparsely labelled *tenm4^+/+^* and *tenm4^-/-^* RGCs, with the same cell imaged at 4dpf and 10dpf. Synaptophysin is labeled with GFP (grey, pointed by white arrowhead) and axonal arbor is labelled with DsRedx (magenta). Scale bar: 10μm. All error bars are mean ± S.E.M. **(D - H)** Quantification of presynaptic puncta volume **(D)**, density **(E)**, and axonal branch length **(F)** in *tenm4^+/+^* and *tenm4^-/-^* RGCs, as well as dot comparison among parameters at 4dpf **(G)** and 10dpf **(H)**, with the Person correlation coefficients r. Each dot represents the data from one zebrafish and n= 34 for each genotype; p-values are from two-way ANOVA repeated comparison without correction and Fisher’s LSD post-hoc test. **(I – L)** 3D reconstruction of *tenm4^+/+^* and *tenm4^-/-^*RGC **(I)** and Sholl analysis of them **(J)** with p-values from genotypes comparison **(K)**, *tenm4^+/+^* vs *tenm4^-/-^*at 4dpf (black curve) and 10dpf (orange curve), and development comparison within genotype **(L)**, *tenm4^+/+^* RGC at 4dpf versus 10dpf (light blue curve); *tenm4^-/-^* RGC at 4dpf versus 10dpf (magenta curve), p-values are from three-way ANOVA repeated comparison with Tukey’s post-hoc test. Scale bar: 10μm.

We first evaluated the potential defects of the *tenm4* deletion on the basic IPL organization and retinal cell numbers between 3 and 5 days post fertilization (dpf), when retinal connectivity is maturing and visual evoked responses emerge. Using *Tg(Isl2b:Gal4;UAS:GFPcaax*) animals, where the membrane-linked GFP is expressed in the majority of RGCs, we found that the typical arrangement of strata in the IPL was not disturbed in *tenm4* mutants compared to the wildtype (WT) pattern and fluorescence intensity measurements did not show any differences (Figure S1C). This suggests *tenm4* is not required for gross IPL stratification of RGCs. Furthermore, phospho- histone H3 (PH3) and terminal deoxynucleotidyl transferase dUTP nick end labeling (TUNEL) assays in the retina showed no obvious changes in cell proliferation and apoptosis, apart from a transient elevation in the number of apoptotic cells at 4 dpf (WT: 85.60 ± 6.91 versus KO: 138.60 ± 10.99, *p < 0.01\*\**; Figure S1D and S1E). Together with the observation of high numbers of Tenm4^+^ neurons in the CMZ region, this could point towards a function in regulating neuron numbers at this developmental stage and therefore influence the overall retinal circuit and output.

To investigate any morphological consequences upon *tenm4* deletion on individual RGCs in the retinotectal projection at 4 dpf, the axonal arbor and presynaptic puncta (as identified by synaptophysin localisation) of RGC were fluorescently marked through sparse cell labelling. To this end, the *ath5:Gal4* and *UAS:Syn-GFPcaax;UAS:DsRedx* plasmids were co-injected into WT animals and *tenm4^-/-^* mutants incross-derived embryos.^44^ This allowed for the reconstruction of cell morphology, such as measuring synaptic puncta size and density from puncta number as well as determining axonal arborization length (Figure 2B, 2C and S2A). We found that the loss of *tenm4* had a strong effect on the volume and density of presynaptic puncta along the RGC arbors in the tectum at 4 dpf, dramatically raising the average volume by 2.4 fold (WT: 0.69 ± 0.09 μm^3^ versus KO: 1.64 ± 0.17 μm^3^, *p < 0.0001\*\*\*\**), and density by 1.3 times (WT: 0.17 ± 0.01 versus KO: 0.22 ± 0.01 puncta/μm, *p < 0.001\*\*\**), while slightly increasing puncta number and reducing branch length (Figure 2D to 2F, and S2A). Consistent with the fact that *tenm4* is not expressed in all RGCs, the enlarged puncta were restricted to a subset of RGCs only, accounting for 21% (7/34) of the population. Furthermore, we found that in tenm4 mutant fish 29% (10/34) of RGC arbors contained some enlarged presynaptic puncta when measured individually and compared to the maximum volume detected in RGCs from WT (Figure S2B). Interestingly, we observed that in the *tenm4^-/-^* mutants RGCs with enlarged puncta were consistently associated with increased puncta density and reduced branch length, showing a significant correlation puncta volume and density (r = 0.4784**, Figure 2G). Moreover, Sholl analysis of reconstructed RGC arbor morphology (Figure 2I) revealed a significant decrease in the number of axonal intersections and thus complexity in *tenm4^-/-^* mutants at 4 dpf (Figure 2J, 2K, and S2D). These data illustrate that loss of *tenm4* leads to an aberrant increase in size of presynaptic structures on RGCs in the optic tectum, possibly due to presynaptic molecules overaccumulating,^45,46^ leading to simplified axonal arbors and synapses clustering on specific branches.

We next explored if these early synaptic defects persist to later developmental stages. For this, we analyzed the presynaptic structures again in the same cells of animals at 10 dpf, when most axonal arborizations are mature (Figure 2C). We found that in *tenm4^-/-^* mutant RGCs, although puncta volumes significantly declined between 4 dpf and 10 dpf (compared to a slight increase seen in WT), the average presynaptic puncta volumes were still significantly increased at the later stage (WT: 0.77 ± 0.10 μm^3^ versus KO: 1.17 ± 0.18 μm^3^, p < 0.05*), indicating a stable effect (Figure 2D and S2C). While puncta density in mutants normalized at this stage compared to WT animals (Figure 2E), axonal branch length was considerably shorter in mutants (Figure 2F). Enlarged puncta were accompanied by shorter branch length, as shown by a significant negative correlation (r = −0.3455*), while the correlation with puncta density at early stages was lost (Figure 2H). Importantly, Sholl analysis revealed that RGCs in *tenm4* mutants are much less complex, exhibiting fewer intersections compared to WT RGCs (Figure 2K and S2E). Additionally, we detected a slowing in progression of structural complexity in mutant animals, as indicated by the Sholl analysis (Figure 2L): significant developmental differences between branch complexity at 4dpf and 10 dpf were observed only in a much smaller range for mutants than the more general effect seen in WT RGCs (Figure S2F and S2G). In summary, *tenm4* loss leads to significant changes in presynaptic structures and their densities, with associated phenotypes in axonal branch formation and complexity.

### Tenm4 mutants show aberrant RGC axonal arbor growth dynamics during early development

So far, we have measured the morphological impact of *tenm4* deletion at defined timepoints of 4 and 10 dpf. To discover whether *tenm4* affects the dynamics of RGC arbor growth, we tracked the innervation of RGCs axons in the optic tectum between 3 and 3.5 dpf using *in vivo* live imaging (Videos S1 and S2). We sparsely labeled RGC axons and the presynaptic marker synaptophysin by co-injection of *ath5:Gal4* and *UAS:Syn-GFPcaax;UAS:DsRedx* plasmids into WT and *tenm4^-/-^*mutant embryos. We first focused on the initial three hours (with 10 min image intervals) when RGCs have arrived in the optic tectum (Figure S3A). Our analysis of branch reconstruction (Figure 3A) revealed that during the gradual increase in branch length and puncta density, RGC axons in *tenm4^-/-^* animals are significantly more dynamic compared to WTs, as indicated by a 2-fold increase in branch motility (3.82 ± 0.28 relative to 1.89 ± 0.14 μm/10 min in WTs, *p < 0.001\*\*\**) and a 1.34-fold increase in puncta turnover (0.023 ± 0.0022 puncta/µm relative to 0.017 ± 0.0013 in WTs, *p = 0.0813*) (Figure 3B to 3E). To further characterize the elevated branch motility, we next focused our analysis on branching events, which are categorized as retracted (present at time-point t_n_ but lost at t_n+1_), transient (lost later >t_n+1_), and newly-added branches (appearing after t_0_ and persisting to the endpoint).^47^

**Figure 3.**
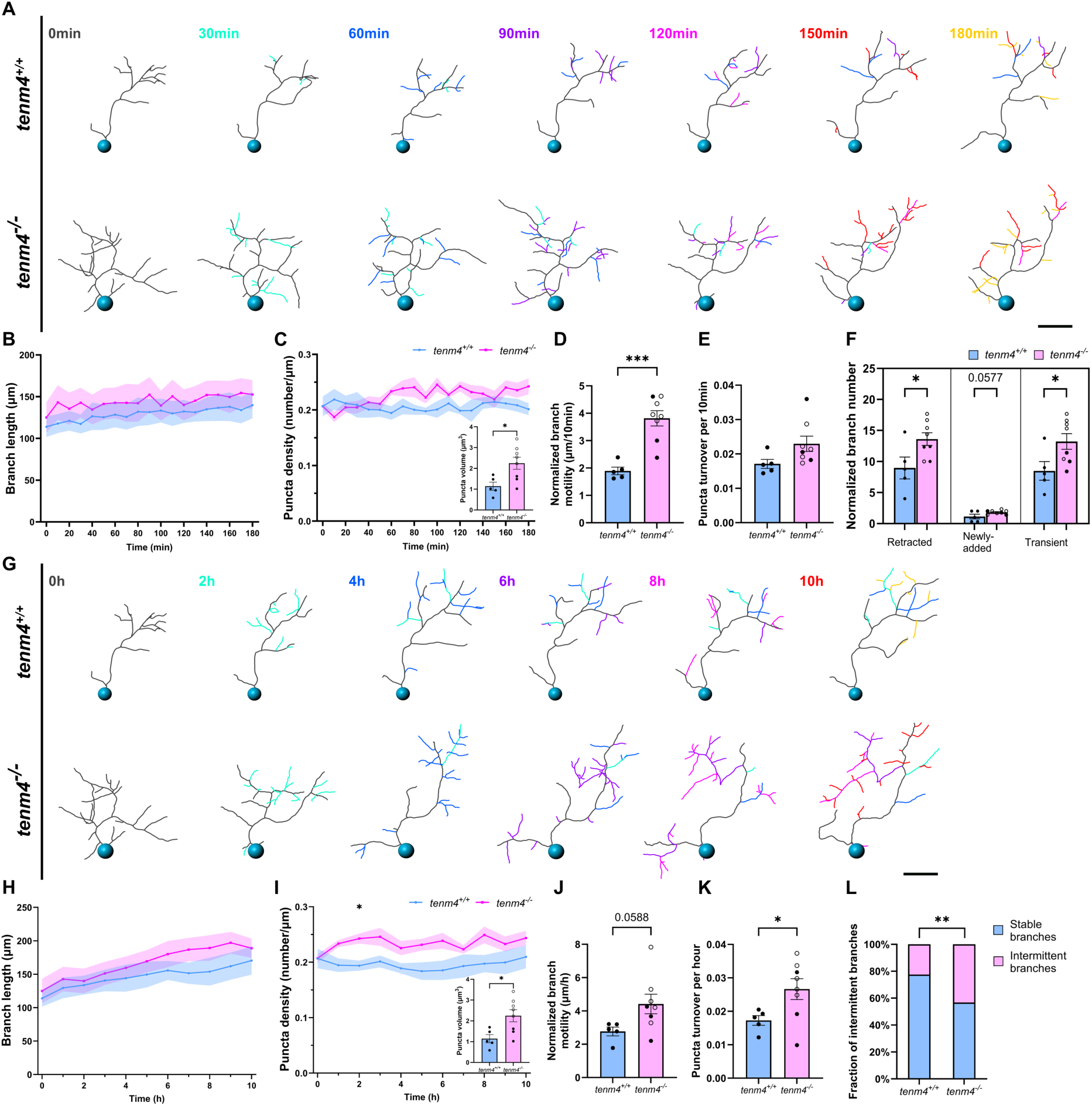
***Tenm4* mutants exhibit increased RGC branch and presynaptic structure dynamics during early development (A and G)** 3D-reconstruction of RGC arborizations from time-lapse images (10 h, 10 min interval) in *tenm4^+/+^* and *tenm4^-/-^* animals from 3 dpf to 3.5 dpf, with color-coded newly formed branches every 30 min during the first 3 h **(A)** and every 2 h between 3 and 10 h **(G).** Blue spheres indicate the position of the first branch point, which was defined as start point for measurements. Scale bar: 10μm. **(B - L)** The growth curve of axonal branch and presynaptic puncta density, as well as normalized branch motility and puncta turnover, indicated by the absolute difference of branch length or puncta density between two consecutive time points (normalized to the number of stable branches), between *tenm4^+/+^*and *tenm4^-/-^* RGCs in the first 3 h with 10 min interval **(B - E)** and 10 h with 1 h interval **(H - K)**. Puncta volume comparison is incorporated. **(F)** The number of branching events including retracted (present at one time-point but lost at the next), transient (lost at a later timepoint), and newly-added branches (appearing at one time-point and persisting to the endpoint) between *tenm4^+/+^* and *tenm4^-/-^*RGCs in the first 3 h with 10 min interval. They are normalized to the number of stable branches (persist over 4 h). **(L)** The fraction of transient branches (initially classified as stable branches but then retracted) in 10 h, indicated by the number of transient branches over the number of stable branches, between *tenm4^+/+^* and *tenm4^-/-^* RGCs in 10 h with 1 h interval. p-values of growth curve are from two-way ANOVA repeated comparison with Šidák post-hoc test, while others are from unpaired t-test. All error bars are mean ± S.E.M. n=5 larvae for each genotype. Hollow circles in *tenm4^-/-^* indicates RGCs with larger puncta size.

In *tenm4^-/-^* mutant RGCs, we found branch addition and elimination events were markedly more frequent, with the number of retracted and transient branches rising 1.52-fold and 1.56-fold, respectively (retracted WT vs KO: 8.97 ± 1.76 and 13.61 ± 1.03, transient WT vs KO: 8.48 ± 1.50 and 13.23 ± 1.26, *p < 0.05*,* Figure 3F). In addition, there was a strong trend for the number of newly added branches showing a 1.61-fold increase (WT: 1.13 ± 0.37 versus KO: 1.819 ± 0.12, *p = 0.0577*). Interestingly, when focusing only on the RGC population with enlarged synaptic puncta in mutant animals (n = 5), we found that the increased branch dynamics were amplified compared to RGCs with normal appearing puncta, again suggesting that in our *tenm4* mutants there is heterogeneity in the phenotypes of individual RGCs (Figure S4A to S4E).

We next assessed branch and synaptic puncta dynamics over a longer period (10 hours, imaged at 1-hour intervals, Figure 3G and S3B). When assessing all RGCs in the *tenm4* mutant animals, we found that, like in the initial 3-hour window, there was a significant increase in puncta turnover (*p<0.05\**) and a strong trend in branch motility (*p = 0.0588*) compared to WT animals (Figure 3J and 3K). When focusing only on RGCs with increased puncta size in mutants (Figure S4F to S4J), the difference in branch motility became statistically significant (*p<0.05\**, Figure S4H). Consistently, branch motility and puncta turnover in this subset of mutant cells were considerably enhanced from 1.6-fold to 1.82-fold and from 1.54-fold to 1.76-fold relative to WT RGCs, respectively (Figure 3J, 3K S4H and S4I). Long-duration imaging enabled us to investigate the turnover rate of stable branches (persisting longer than 4 hours). We found that mutant RGC branches were remarkably unstable (Figure 3G), with a greater proportion of them previously classified as stable branches then transformed into transient branches in this time window (44.12% versus 22.5% in WT, Figure 3L and S4J).

In summary, our long-term morphological analysis illustrates that *tenm4* is required for correct axonogenesis and synaptogenesis of RGC from the initial stages of axon ingrowth into the target, the optic tectum. Loss of *tenm4* leads to significant changes in branch formation and dynamics, as well as the generation and turnover of synaptic puncta, suggesting a critical role for this protein in the establishment of functional connectivity in the visual system.

### Stronger functional response of RGCs upon Tenm4 deletion

Given the morphological synaptic aberrancies we detected in *tenm4* mutant larvae, we next investigated the possible functional consequences. First, we assessed if the aberrant presynaptic puncta of individual RGCs are indeed functional and release glutamate. For this we expressed a fluorescent glutamate reporter sparsely in RGCs through injections of ath5:Gal4 and UAS:SFiGluSnFR plasmids at the one cell stage.^48,49^ At 4 dpf we then presented drifting bars with 100% contrast to one eye while functionally imaging individual RGC presynaptic puncta in the contralateral tectum (Figure 4A). We detected a significant 2-fold increase of stimulus-evoked glutamate release in RGCs from *tenm4* mutants compared to WT animals, when assessing the average peak amplitude (Figure 4B). This phenotype was accompanied by a prolonged overall response (ΔF/F) and a significantly longer decay (3.81-fold) back to baseline fluorescence, in comparison to the rapid and transient response seen in the WT group (WT: 0.335 ± 0.022 s versus KO: 1.276 ± 0.137 s, *p < 0.0001\*\*\*\**; Figure 4F). Interestingly, the peak amplitude of individual events remained unchanged between the two groups (Figure 4C), suggesting that the aberrant increase in mutant RGCs was driven by a more sustained response. This was further reflected by a significantly higher cumulative response (WT: 5.420 ± 0.247 versus KO: 9.735 ± 0.355, *p<0.0001\*\*\**, Figure 4D), as indicated by the area under the curve (AUC) measured above the baseline (ΔF/F = 0), and a significantly lower total number of individual responses (WT: 24.347 ± 0.565 versus KO: 20.441 ± 0.214, *p<0.0001\*\*\*\**, Figure 4E), respectively. These phenotypes were also evident in the relative frequency distribution of glutamatergic puncta, where a greater proportion of presynaptic sites in mutant RGCs exhibited larger glutamate release events, associated with fewer response frequencies and slower signal decay, while response latency remained largely unaffected (Figure S5). In summary, the aberrant presynaptic structures are fully functional and loss of *tenm4* results in a lower frequency of events, but an increased, sustained, and poorly regulated efflux of glutamate from individual pre-synaptic RGC boutons.

**Figure 4.**
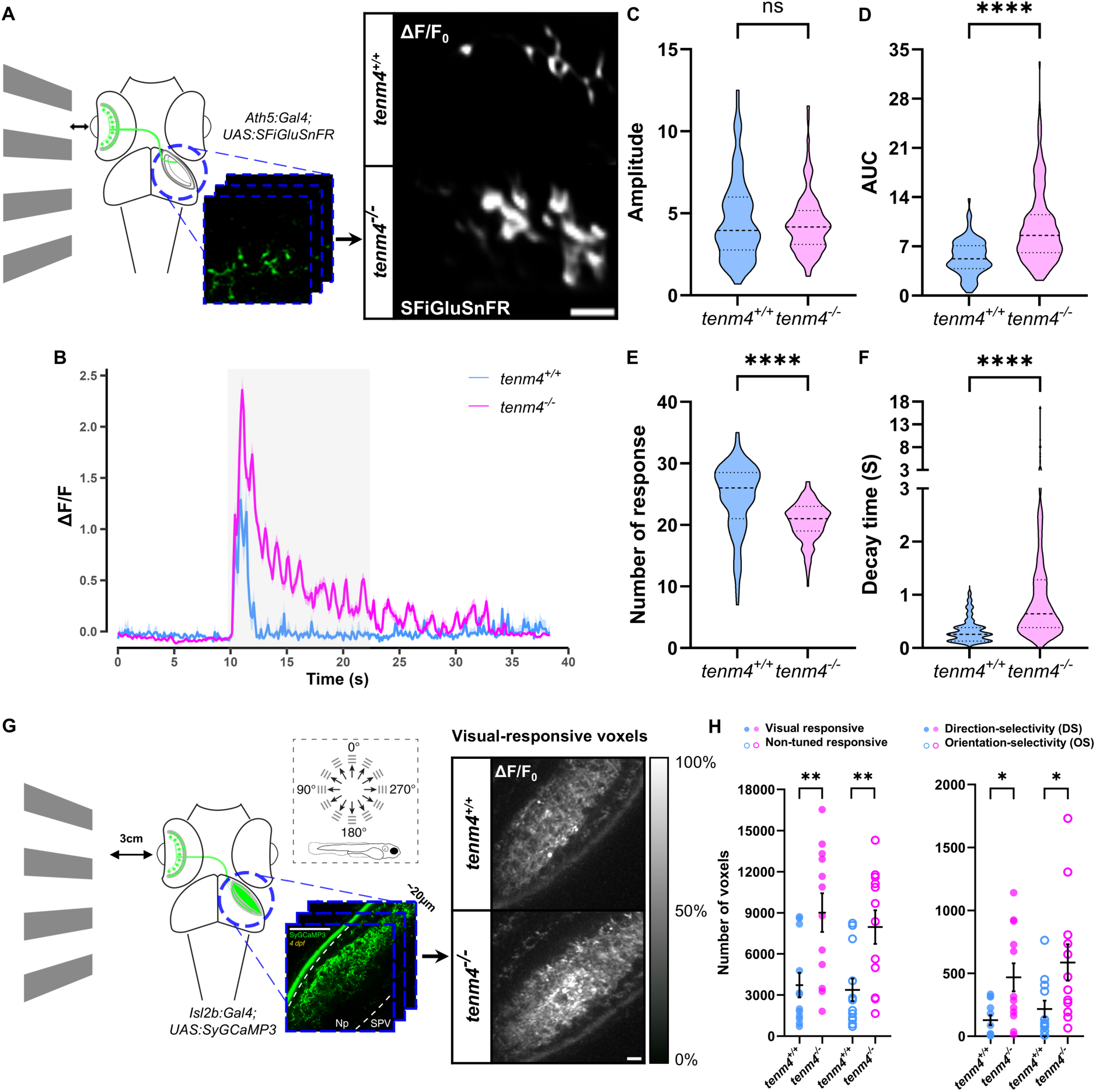
Stronger functional response of RGCs upon *tenm4* deletion (A and. **B)** In vivo set up for imaging glutamate release by individual RGC axon terminal in 4dpf *tenm4^+/+^* and *tenm4^-/-^* larvae, labelled by co-injection of ath5:Gal4 and UAS:SFiGluSnFR **(A)**. Stimulus-evoked glutamate response in two genotypes is shown by the average ΔF/F_0_ over 300-time frames (∼40 s) **(B)**. Scale bar: 10μm. **(C to F)** The amplitude-peak response **(C)**, area under curve above the baseline (y=0)- cumulative response **(D)**, total number of response **(E)**, and decay time **(F)** were measured from individual region of interest (ROI, indicating glutamate puncta); n=101 ROIs for *tenm4^+/+^* and n=220 ROIs for *tenm4^-/-^,* and p-values are from unpaired nonparametric two-tailed t-test (Mann-Whitney test). **(G)** In vivo functional calcium imaging set up for detecting SyGCaMP3 (green) expressed by RGC axon terminals in 4dpf *Tg(isl2b:Gal4;UAS:SyGCaMP3)* larvae. Moving bars at different angles relative to larval orientation (showing in the dashed box) is projected to the screen (3cm away from the larvae). Response is recorded at 2-4 Z-planes around the center of the contralateral optic tectum, with mean ΔF/F_0_ calculated in *tenm4^+/+^* and *tenm4^-/-^*. Scale bar: 10μm. Np: neuropil; SPV: stratum periventriculare. **(H)** Average number visually responsive and non-tuned voxels, as well as direction- and orientation-selective voxels in *tenm4^+/+^* and *tenm4^-/-^*; n=12 larvae and p-values are from unpaired nonparametric two-tailed t-test (Mann-Whitney test).

To investigate the functional responses of RGCs at the population level, we analyzed calcium responses of RGC axon terminals innervating the tectal neuropil in animals expressing the genetically encoded calcium indicator GCaMP3 fused to synaptophysin in all RGCs. Drifting bars moving in 12 directions were presented to one eye of 4 dpf Tg(*Isl2b:Gal4;UAS:SyGCaMP3*) transgenic larvae while functionally imaging the contralateral tectum (Figure 4F), as previously described.^50^ Visually responsive voxels were isolated by voxel-wise analysis. *In vivo* calcium imaging revealed that *tenm4* deletion induced generally stronger responses from RGC axonal terminals at 4 dpf, largely increasing the total number of voxels (WT: 3,724 ± 892.2 versus KO: 9,014 ± 1,412 pixels, *p<0.01\*\**), with about 50% of mutant animals showing extremely high response (Figure 4G). Further analysis showed that this increase in *tenm4* mutant animals was present in direction- and orientation- selective (DS and OS, 127.4 ± 39.68 versus 468.1 ± 111.8 pixels, 216.7 ± 65.96 versus 586.3 ± 144.9 pixels in WT and KO, respectively, *p<0.05\**) and non-tuned responses (non- DS and OS, WT: 3,380 ± 819.1 versus KO: 7,960 ± 1,224 pixels, *p<0.01\*\**), Figure 4H). These findings suggest that the effect is not RGC subpopulation-specific, in line with our expectations based on the broad *tenm4* expression in the retina.

### Visually-driven behaviors are impaired asymmetrically in Tenm4 mutants

Since *tenm4* was highly expressed in the retinorecipient AF 6-9 (Figure 1E), which includes the pretectum essential for inducing looming-evoked and optic flow-dependent visuomotor behaviors, we next evaluated the ramifications of the *tenm4* deletion for visual behaviors. For this we focused on the optokinetic response (OKR) and the optomotor response (OMR). The OKR and OMR are visually-induced compensatory eye or body movements to stabilize gaze or position to limit retinal image displacement.^51^ In the OKR, immobilized animals perform eye saccades along the direction of moving stripes presented along the nasal and temporal axis (Figure 5A).^52^ During OMR, animals perform locomotor responses, aligning their body movement with perceived translational motion.^53^ Therefore, when presented with grated bars moving perpendicular to the zebrafish larval body axis in left or right directions, the animals will make counterclockwise or clockwise turns, respectively (Figure 5G).

**Figure 5.**
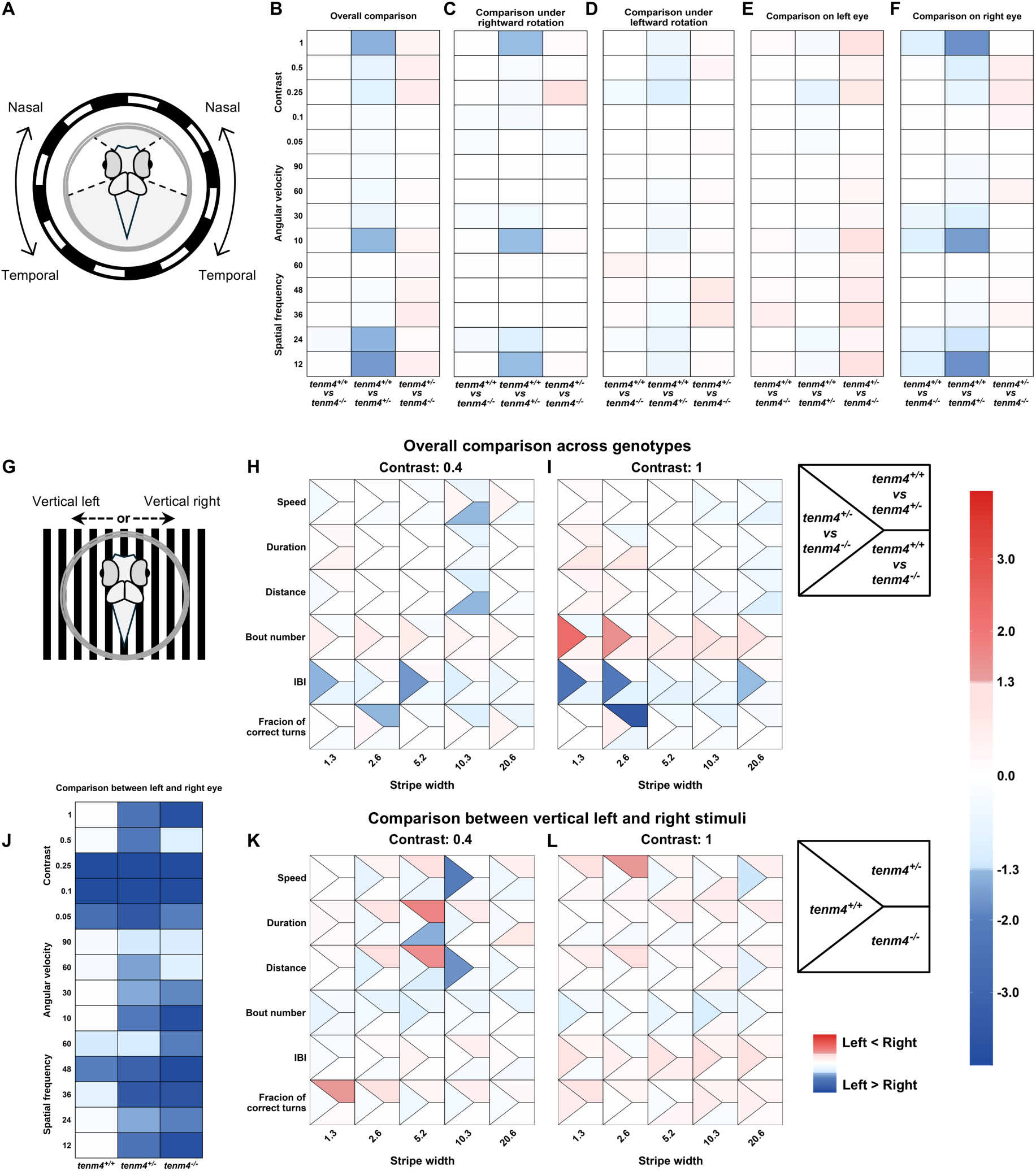
Visual acuity assay reveals left-right asymmetrical impairment in *tenm4* mutants (A and. **G)** Schematic showing OKR (A) and OMR (G) set up for the 5 dpf larvae derived from incrossed *tenm4^+/-^*heterozygous animals. Rotating stimuli between nasal and temporal directions are provided to OKR test, while translation stimuli in either leftward or rightward direction perpendicular to fish’s orientation are provided to test optomotor responses. **(B - F)** Heat map of p-values showing the comparison of OKR data among three genotypes in different stimuli varying in spatial frequencies, angular velocities and contrast. The different columns depict *tenm4^+/+^* versus *tenm4^-/-^*, *tenm4^+/+^*versus *tenm4^+/-^*, *tenm4^+/-^* versus *tenm4^-/-^* comparisons. These include overall comparison **(B)**, comparison under rightward rotation **(C)** and leftward rotation **(D)**, comparison on left **(E)** or right **(F)** eye’s sensitivity; n=12 larvae for *tenm4^+/+^* and *tenm4^-/-^*, n=28 larvae for *tenm4^+/-^.* **(H, I)** Heat map of p-values showing the overall comparison of OMR test among three genotypes in different stimuli varying in stripe width and contrast at 0.4 **(H)** and 1 **(I)**. Each cell is divided into a left triangle (*tenm4^+/+^* versus *tenm4^-/-^*), an upper right trapezoid (*tenm4^+/+^* versus *tenm4^+/-^*) and a bottom left trapezoid (*tenm4^+/-^*versus *tenm4^-/-^*). For heatmaps (B - I): A blue color indicates the performance of *tenm4^+/+^* is better than any of the mutants (trapezoids) or the *tenm4^+/-^* is better than the *tenm4^-/-^*(triangle). A red color indicates the opposite. n=34 larvae for *tenm4^+/+^*, n=28 larvae for *tenm4^-/-^*, n=54 larvae for *tenm4^+/-^.* **(J - L)** Heat map of p-values showing the comparison of OKR sensitivity between left and right eye **(J)** as well as OMR activity under vertical left stimuli and right condition at contrast 0.4 **(K)** or 1 **(L)** within the same genotype, where each cell is divided into a left triangle (*tenm4^+/+^*), an upper right trapezoid (*tenm4^+/-^*) and a bottom left trapezoid (*tenm4^-/-^*). For heatmaps (J - L): A blue color indicates the sensitivity in the left eye is better than in the right eye, or the performance under vertical left stimuli is better than under vertical right stimuli. The red color indicates the opposite. All p-values for genotypes comparison are obtained from two-way ANOVA repeated comparison with Tukey post-hoc test, while p-values for lateralized comparison are obtained from three-way ANOVA repeated comparison with Tukey post-hoc test.

The OKR and OMR emerge in zebrafish larvae from 3 dpf and 5 dpf, respectively^51^. We therefore measured behavioral phenotypes in larvae from *tenm4^+/-^* heterozygous (Het) incrosses at 5 dpf blind to genotype, with post-experimental genotyping. For the OKR, we quantified the slow phase of optokinetic nystagmus, as indicated by gain, which is the ratio of eye velocity to stimulus velocity, thereby sensitivity to the stimuli, across grating with different spatial frequencies (SF), angular velocity (V), and contrast. No changes were observed in *tenm4^-/-^* mutants regardless of whether the stimulus rotated in either leftward or rightward direction (Figure 5B to 5D, S6A to S6E). However, when splitting the data between the two eyes, we noticed an attenuated response in the right eye towards rightward direction in *tenm4^+/-^* mutants under multiple conditions (SF = 12 or 24 cycles / 360 degrees, V = 10 degrees / second and contrast = 1), where the gain was robustly reduced by approx. 13% compared to the left eye (*p<0.05\**, Figure 5B to 5F, S6A to S6G). This data suggests that *tenm4* controls visual perception asymmetrically. We therefore compared interocular responses under whole-field motion. While we found lateralized visual sensitivity in WT larvae under several extreme challenging conditions with low contrast, both the *tenm4* hetero- and homozygous mutants exhibited significantly enhanced phenotypic lateralization in virtually all cases, including in normal conditions (Figure 5J and S6F, Table S2). Responses from the right eye were remarkably slower than left-eye responses. Such a left-right phenotypic bias in *tenm4* mutants is consistent with the asymmetric tenm4 expression we detected earlier, showing higher *tenm4* levels present in the right eye (Figure 1F). Importantly, we did not detect any differences in oculomotor performance (position and velocity) across genotypes (Figure S6H and S6I), suggesting that *tenm4* does not influence general ocular movement and extraocular muscle function.

We next tested the visual performance in the OMR, varying stripe width (SW) and contrast (Figure 5G to 4I).^54^ In the vast majority of conditions, *tenm4* mutant larvae did not show a general locomotor dysfunction with respect to swimming speed, distance and bout duration, except at low contrast condition (0.4) and a SW of 10.3 μm. In this condition, *tenm4^-/-^* mutants moved slower (6.82%) and covered a shorter distance (10.43%) compared to WT larvae (Figure 5H, 5I and S7). We further found that *tenm4^−/−^*mutants displayed a hyperactivity phenotype (increased bout numbers, BN; decreased interbout interval, IBI), whereas *tenm4^+/-^* larvae were less active, exhibiting fewer BNs and increased IBI for some conditions. Consequently, significant changes were observed between the two mutant genotypes at SW of 1.3 μm (*p<0.05\**) and 5.2 μm (*p<0.01*\*\*) under a low-contrast condition for IBI, and at SW of 1.3 μm (*p<0.01*\*\* for BNs and IBI), 2.6 μm (*p<0.05*\* for BNs and *p<0.01*\*\* for IBI), and 20.6 μm (*p<0.05*\* for IBI) under optimal contrast (Figure 5H, 5I and S7). Moreover, *tenm4^+/-^*larvae were more prone to making incorrect turns under both optimal and low contrast contrast conditions at a SW of 2.6 μm, with a difference of 10.7% (*p<0.001*\*\*\*) and 6.4% (*p<0.05*\*), respectively, relative to WT larvae (Figure 5H, 5I and S7).

These phenotypes primarily resulted from vertical left stimuli rather than vertical right stimuli, which led to variation in IBI (Figure S8). We therefore examined laterality during OMR under left and right direction of stimuli (Figure 5K, 5L and S8). Although WT larvae showed a slight leftward bias in bout speed and distance under a more challenging condition (width=10.3 μm, contrast=0.4), we found a clear lateralization of phenotypes in *tenm4* mutants even under more favorable conditions (Figure 5J), in which Hom larvae exhibited longer bout duration during left stimulation (width = 5.2 μm, contrast = 0.4) whereas Het larvae were more active during right stimulation, with longer bout duration and distance (width = 5.2 μm, contrast = 0.4), as well as a higher fraction of correct turns (width = 1.3 μm, contrast = 0.4).

Taken together, these data demonstrate that the loss of the asymmetrically expressed *tenm4* leads to lateralized defects in OKR and OMR performance. Furthermore, we find that deficiencies in visual perception are more pronounced in heterozygous animals than in homozygous animals, possibly suggesting dominant-negative mechanism.^55,56^

## DISCUSSION

Comprehensive studies have provided compelling evidence that teneurins, as synaptic adhesion molecules, contribute to mechanisms underlying neural circuit assembly.^15,31,57,58^ However, our full understanding of their roles in vertebrate neural circuit wiring, including the visual system, is still incomplete. Here, we showed that Tenm4 is more broadly expressed across the visual system, including all types of retinal cells and TCs. Within the retina, Tenm4 expression was particularly enriched in the ciliary marginal zone (CMZ), the principal site of retinal neurogenesis, suggesting that Tenm4 may contribute to retinal cell differentiation during development. This interpretation is consistent with the established roles of teneurins in regulating cellular differentiation in other tissues, including human TENM2 during odontoblast development^59^ and adipocyte differentiation^60^, and mouse Tenm4 during oligodendrocyte differentiation^28^.

Interestingly, we found that Tenm4^+^ RGCs and their main postsynaptic cellular targets TCs are more abundant on the right side of the animal compared to the left side, suggesting a lateralized role in the visual system. Consistent with this hypothesis, when we examined visual behavioral in *tenm4*, we observed a phenotype specifically in the right eye in both *tenm4^+/-^* Het and *tenm4^-/-^*Hom mutant larvae during OKR, and mutants also showed locomotor lateralization between vertical left and right stimuli under non-challenging visual conditions during OMR. Recent studies show that OKR and OMR under global motion are mostly symmetric.^55,56^ However, based on monocular stimulation experiments, there is evidence that the sensorimotor circuits in zebrafish are not completely independent between the two animal sides but instead are differentially organized. There is coordination and processing of information across the hemispheres through the posterior commissure, the tectal commissure and the hindbrain, for example in response to gratings from different spatial locations, directions, and orientations.^56,66^ Such processing organization has also been shown in the small-field motion-like prey capture, where interhemispheric communication is mediated by GABAergic inhibitory neurons through commissural tectal circuitry that traverses the tectal commissure.^67^ Notably, not only the retinotectal circuit, but also in the pretectum, the first optic-flow processing center,^68,69^ and the reciprocal network of other tectal targets including thalamus, nucleus isthmi, torus longitudinalis, the nucleus of the medial longitudinal fasciculus as well as cerebellum and medulla oblongata in hindbrain containing ventromedial spinal projection neurons mainly constitute the sensorimotor circuit.^41,66,70^ In these regions, as well as in regions involved in interhemispheric communication, Tenm4 exhibits moderate expression, suggesting involvement in the construction of visual circuits and visual behavioral phenotypes, which is consistent with our observed OKR and OMR phenotypes in *tenm4* mutant animals. The stronger dysfunction originating from heterozygous suggests a dominant-negative mechanism in this genotype or genetic compensation on homozygous animals, whereby the coexistence of WT and mutant proteins could lead to the formation of aberrant protein complexes that alter signaling transduction and behavioral ouputs.^71,72^ This heterozygous effect has also been demonstrated in Tenm4-related neural disorders.^24,73–75^

RGCs are the only retinal output and therefore deliver all perceived visual signals to other brain regions. We present evidence that *Tenm4* is required for correct structural and synaptic development of RGCs in zebrafish, with a role from as early as axons target the optic tectum around 3 dpf. The growth dynamics and complexity of RGC axonal remodeling are both affected by the loss of *tenm4*. The accumulation of synaptic proteins during circuit development is highly correlated with axonal branch formation. ^44^ Importantly, we find that in Tenm4 mutants we detect aberrant synaptophysin localization and accumulation along axonal branches. Changes in presynaptic structure and size upon gene deletions have been clearly reported in *Drosophila* at the neuromuscular junction, with responsible genes regulating synaptic growth, cytoskeletal processes, cell adhesion, vesicle trafficking or BMP/Wnt signaling.^76,77^ In vertebrates there are fewer examples, however the molecules belong to similar categories, including active zone organization.^78^ Although Tenm4 has been linked to cytoskeletal organization,^79,80^ it is currently not known if this molecule also interacts with active zone proteins. However, it seems clear that mislocalization of synaptic proteins can impact on branching behavior in axons. Indeed, we find enlarged synaptic puncta upon *tenm4* deletion that are linked with higher puncta density at early stages and shorter branch length at late stages, suggesting that *tenm4* balances receptor accumulation and synaptic distribution to control synaptic activity and axonal pruning. We furthermore detect significant changes in branch dynamics during the establishment of the retinotectal projection, when RGCs are growing into the tectum and elaborate their arborizations. This could occur directly by regulating synaptic puncta (size and density) thereby ensuring synaptic specificity, or by modulating activity. Alternatively, it may act implicitly through cytoskeletal regulation^79,80^ or the mediation of oligodendrocytes differentiation^29,81^, as oligodendrocyte precursor cells are known to influence RGC outgrowth.^82^ This phenotype is also consistent with stronger response trigger by loss of *tenm4*, which could be associated with a potential role in the modulation of NMDA and AMPA receptor^83^ as well as glutamate recycling at the synapse^84^. Since the morphological phenotype slightly diminishes at a later stage, a compensatory mechanism underlying homeostatic synaptic scaling may be triggered to consolidate the synaptic contact during development.^85^ Interestingly, our observed phenotype seems to be restricted to a subset of RGCs in *tenm4* mutant animals. This can be explained by the fact that *tenm4* is not expressed in all RGCs and therefore a loss of this protein would only affect the normally Tenm4 -positive neurons. However, it is possible that additional RGC subtypes (i.e. normally *tenm4*-negative RGCs) might be affected by *tenm4* loss as well, due to possible heterophilic molecular interactions with tectal cells that have lost Tenm4 on the postsynaptic side.^42,43^ Further experiments are needed to determine exact protein localization across the synapses, either pre- or post-synaptic or both. Additionally, transsynaptic mapping of Tenm4^+^ pre- and post-synaptic site would provide insights into the synaptic interactions underlying circuit function,^86^ while simultaneously enabling the assessment of effects on postsynaptic terminals. Crucially, it will be important to identify the intracellular molecular interactors for Tenm4 to better understand its role in synapse assembly and localization. In summary, our findings support the vital role of teneurins during neural circuit development in vertebrates, including morphological and behavioral consequences.

## ACKNOWLEDGMENTS

This work was supported by grants to RH from the Biotechnology and Biological Sciences Research Council (BB/M000664/1 and BB/R000972/1) and the Medical Research Council (MR/N026063/1 and MR/W006251/1), a PhD fellowship from the King’s-China Scholarship Council (CSC) to ZZ and a PhD Scholarship from the Consejo Nacional de Ciencia y Tecnología, Mexico (CONACYT) to AS-S. The zebrafish central facility at King’s College London (KCL) are acknowledged for their help with animal care. We thank Steve Wilson, Martin Meyer, and Takeshi Yoshimatsu for providing expression plasmids; Phoebe Reynolds and Isaac Bianco for help with the OMR and OKR set ups, respectively; Yan to Ling for input on image analysis; and members of the CDN team for insightful discussions.

## AUTHOR CONTRIBUTIONS

RH conceived the overall study and with help of ZZ, designed the experiments. ZZ created the Gal4 insertion transgenic fish line, performed confocal imaging (expression and morphology), time-lapse imaging and behavioral studies. AS performed basic retinal cellular homeostasis studies and pilot morphological studies. LD performed the functional imaging experiments using the glutamate sensor, while KT performed the functional calcium imaging study. ZZ and RH wrote the manuscript with input from other authors.

## DECLARATION OF INTERESTS

We declare no competing interests.

## MATERIAL AND METHODS

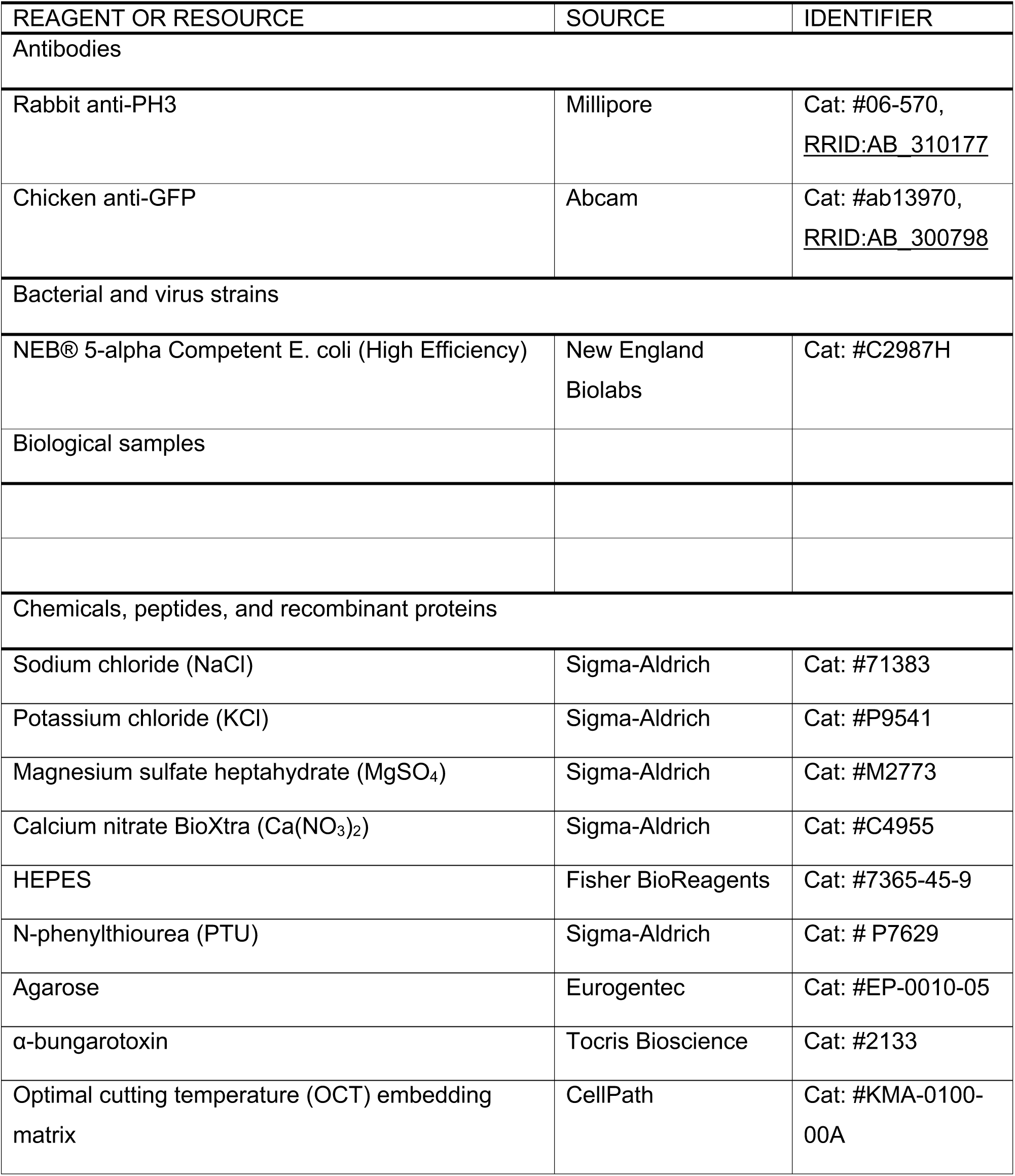
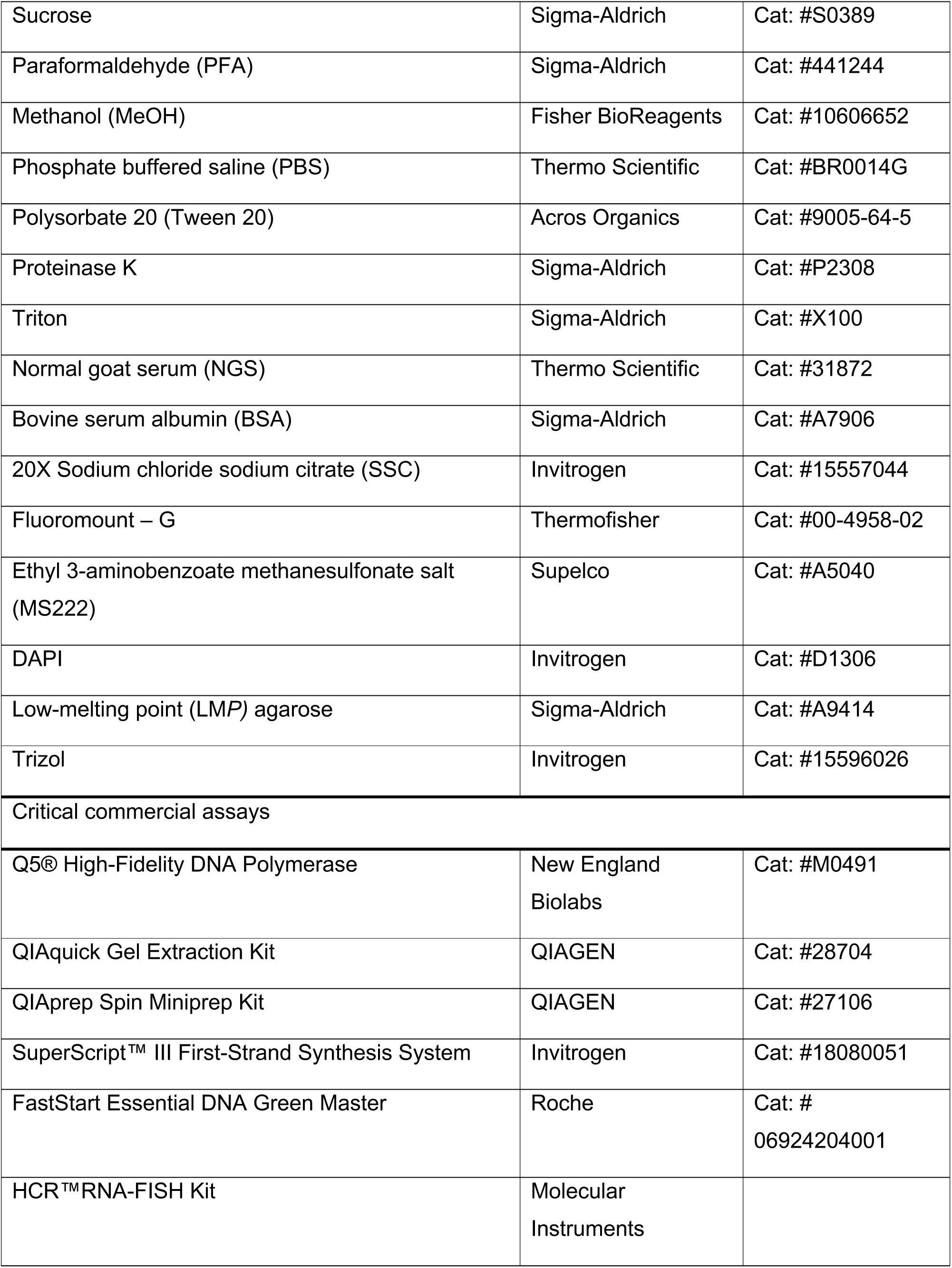
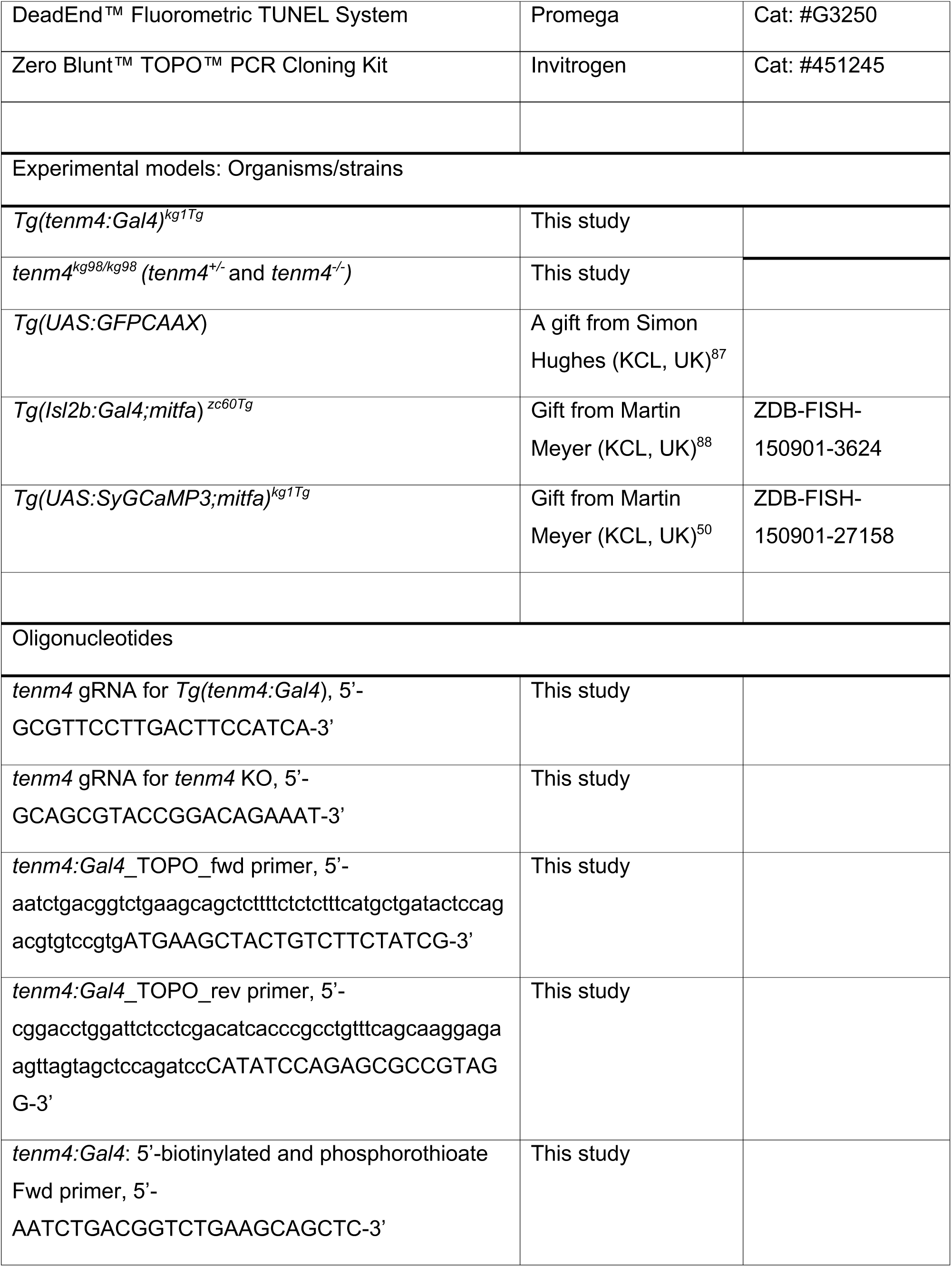
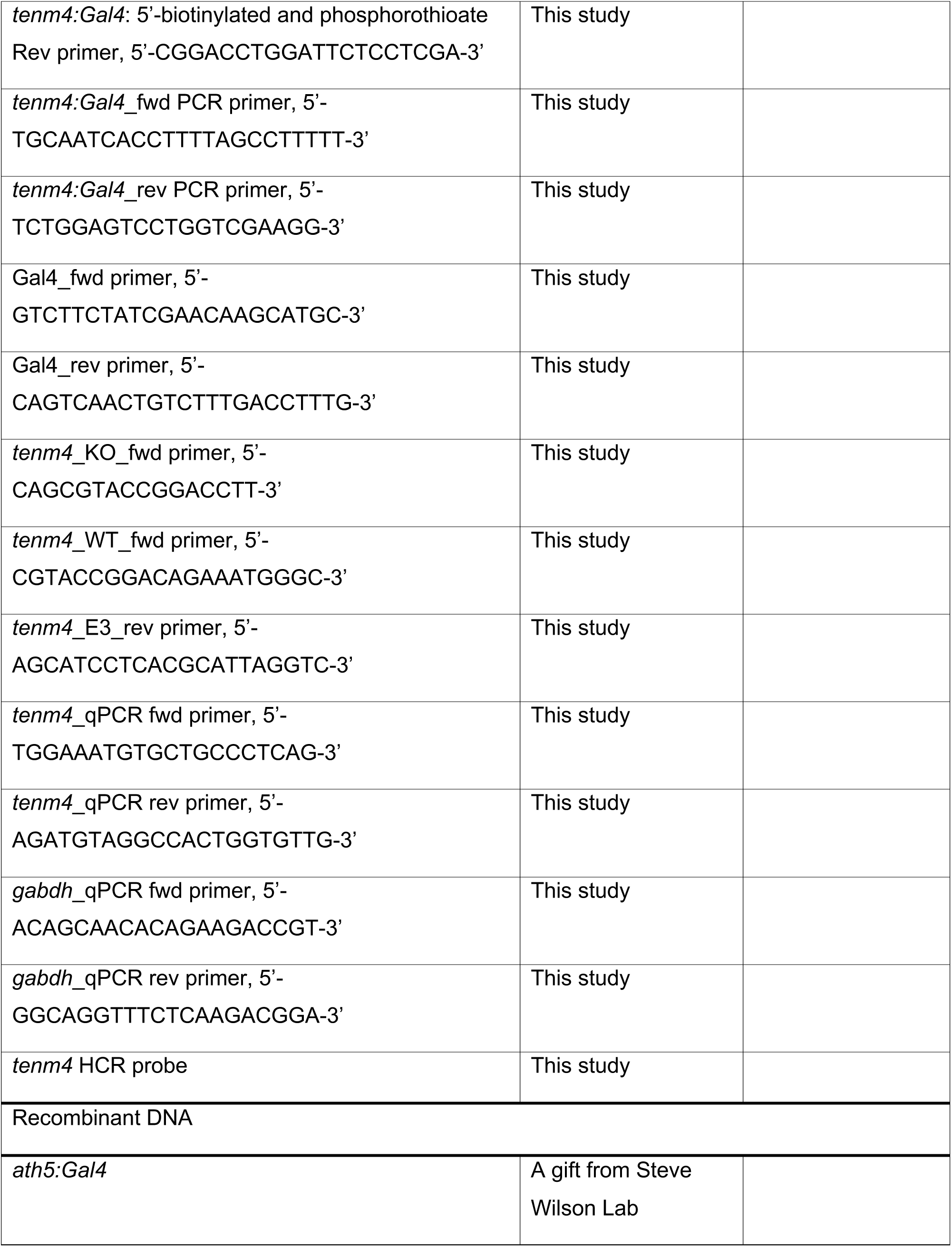
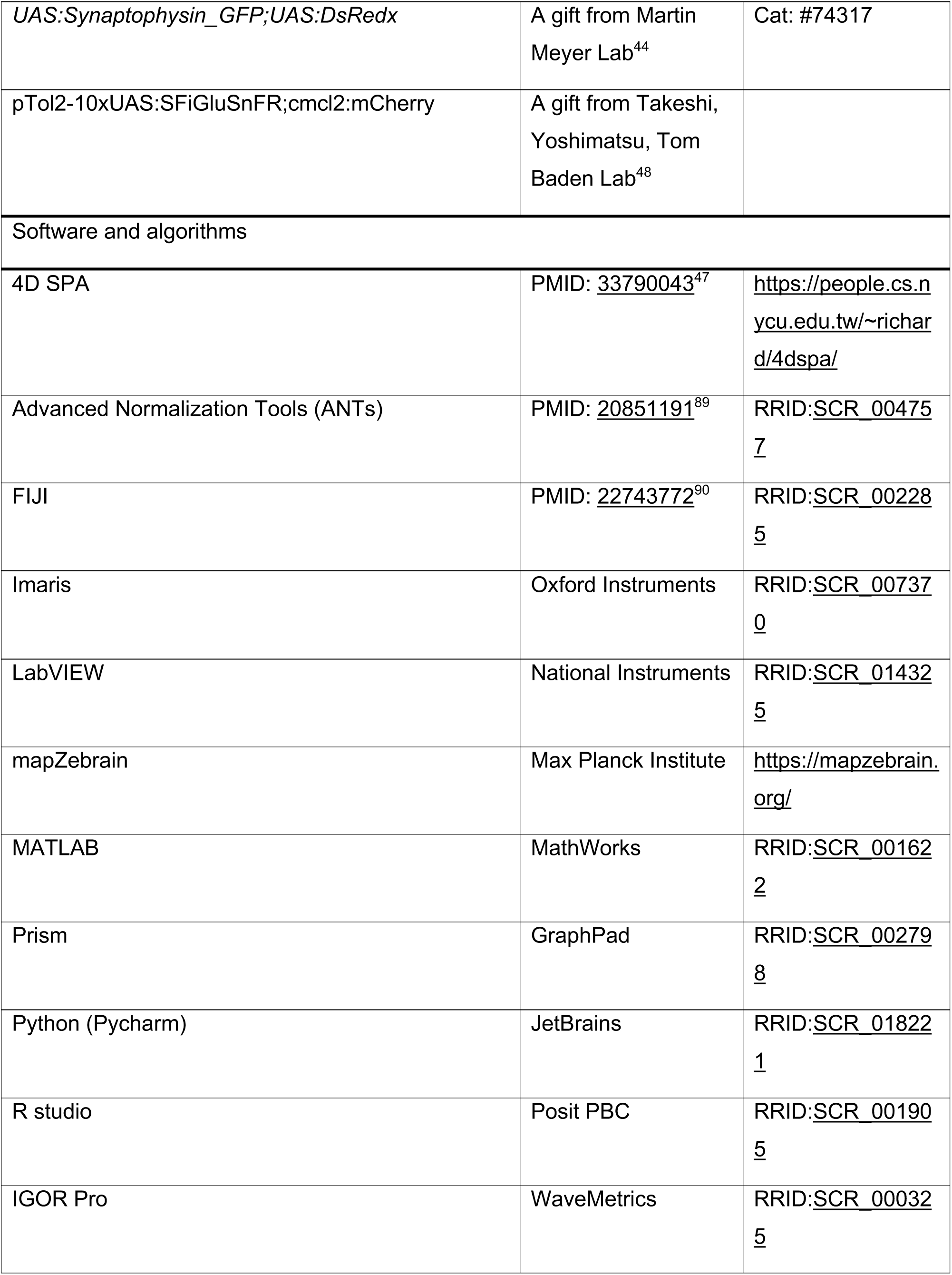
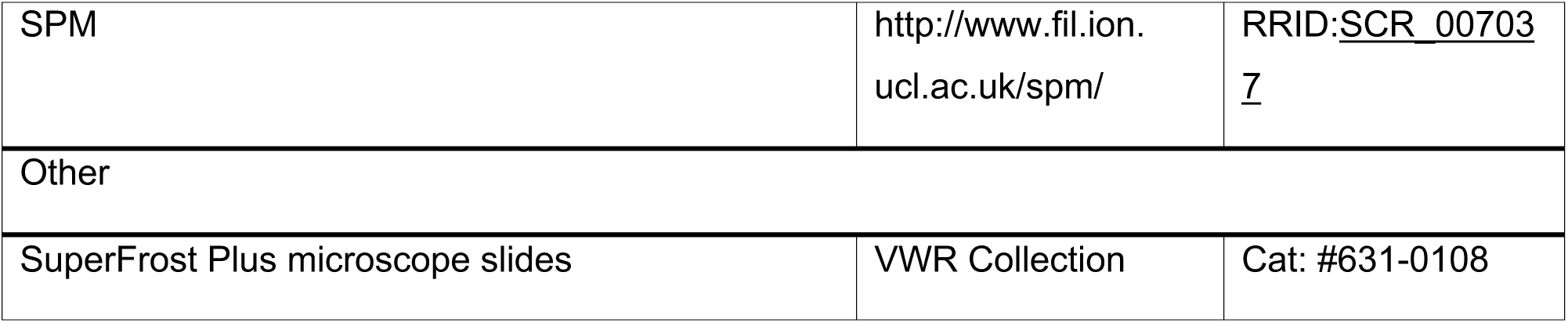

### EXPERIMENTAL MODEL AND STUDY PARTICIPANT DETAILS

Zebrafish husbandry and experiments were carried out in accordance with the UK Animals (Scientific Procedures) Act 1986 and approved by the Animal Welfare and Ethics Review Body at KCL, with zebrafish taken care of by the Zebrafish Central Facility at KCL. All zebrafish experiments were conducted under the Home Office project license PP7266180 to R.H. Embryos were cultured in 90mm Petri dishes (density < 50) containing Danieau solution [58 mM NaCl, 0.7 mM KCl, 0.4 mM MgSO_4_, 0.6 mM Ca(NO_3_)_2_, 5.0 mM HEPES, pH adjusted to 7.6] (additional 200 μM N-phenylthiourea (PTU) was added for structural imaging). Embryos for OKR and OMR were cultured in filtered fish facility water. They were placed in an incubator at 28.5 °C on a 14-hour ON/10-hour OFF light cycle. For experiments on larvae over 5 days, larvae were fed live rotifers according to the KCL Fish Facility feeding protocol (1.5 ml rotifer solution per 5 fish). Structural imaging experiments were performed on the transgenic line *Tg(tenm4:Gal4*) crossing with *Tg(UAS:GFPCAAX*), and AB WT and *tenm4^-/-^* mutants, while behavioral experiments were conducted on the *tenm4^+/-^* Het-incrossed derived embryos. All experimental embryos were aged from 3 dpf to 10 dpf, at which point sex cannot be identified, and were anaesthetized in 1.5 mM MS222 prior to experiments.

### Generation of transgenic lines

The 10 bp knock-out *tenm4^-/-^* mutants line and knock-in line *Tg(tenm4:Gal4*) were generated using CRISPR-Cas9 techniques. The sequence to target the *tenm4* open reading frame (ENSDART00000175825.2) in zebrafish (GRZ11) was selected using CHOPCHOP (https://chopchop.cbu.uib.no/).

The crRNA sequence to create the *tenm4^-/-^* line was designed to target exon 3 on the reverse strand. 0.5 μL of 100 μM crRNA and tracrRNA were mixed at 95 °C for 5 mins to make a 3 µl 3 µM gRNA solution. It was then mixed with 3 μL of 0.5 µg/µL Cas9 protein at 37 °C for 10 mins to assemble CRISPR ribonucleoprotein (RN*P)* complex. F_0_ founders were created by injecting 1 nL of the solution containing 6μL of RNP complex and 0.6 μL of phenol red into AB WT embryos at the one cell stage. Genotyping was used *tenm4*_KO_fwd primer and *tenm4*_E3_rev primer to identify Homs and Hets, while using *tenm4*_WT_fwd primer and *tenm4*_E3_rev primer to identify WTs and Hets. Generation of *Tg(tenm4:Gal4*) followed the established protocol.^34^ Briefly, the *Gal4-VP16-P2A* fragment with *tenm4* 5’ and 3’ homologous arms was PCR amplified using forward (fwd) primer with additional *tenm4* 5’UTR (62 bp) and reverse (rev) primer with additional 66 bp of the downstream sequence of start codon. It was then cloned into pCR 4-Blunt TOPO vector (ThermoFisher) to make a template in the following PCR amplification to generate homology-directed repair (HDR) fragment using 5’-biotinylated and phosphorothioate fwd and rev primers of 20 bp (IDT). All PCR were performed using Q5 polymerase (NEB) with 35 cycles in a final volume of 50-100 µl, in which products were electrophoresed on 1% agarose gels and purified using QIAquick Gel Extraction Kit (QIAGEN) with final elution of 30 µl of warm nuclease-free water, according to the supplier’s instructions. The crRNA sequence was designed to target the exon 2 around the start codon. RNP complex was produced as described above, with 2.5 µl of 10 µM gRNA combining with 0.5 µl of 61 µM Cas9 protein, then mixed with 80 ng HDR template, 10% Cas9 working buffer (20 mM HEPES, 150 mM hydrochloric acid, pH 7.5.) and 10% phenol red at final volume of 5.5 µl for microinjection, as described above.

Cleavage and integration efficiency were assessed in several pools of five injected embryos at 4 dpf using PCR. Germline transmission was screened by Sanger sequencing and fluorescent signals, and one 10 bp KO fish and positive knock-in fish were raised to the next generation. All primers and crRNA sequences were listed in the Key Resource Table.

#### Whole-mount hybridization chain reaction fluorescence in situ hybridization (HCR-FISH)

HCR experiments were conducted on the transgenic knock-in larvae *Tg(tenm4:Gal4;UAS:GFPCAAX*) at 6 dpf. Probes including *tenm4* and *gad1b* were designed using previously published protocol.^91^ All HCR probes and reagents were purchased from Molecular Instruments, and staining was according to the supplier’s instructions. The larvae were fixed overnight at 4 °C in ice-cold 4% PFA/PBS with gentle shaking, then washed 3 x 5 min in PBS to stop fixation, followed by several 10 min treatments in 100% MeOH for dehydration and permeabilization. They were kept at −20 °C overnight or directly transferred 20 larvae for rehydration with a gradient of MeOH/0.1%PBS-Tween (PBSTw): 75% MeOH/25%PBST2, 50% MeOH/50%PBSTw, 25% MeOH/75%PBSTw, 5 ×100%PBSTw for 5 min each at room temperature (RT). The rehydrated larvae were treated with proteinase K (30 µg/mL) for 50 min and washed 2 x 5 min in PBSTw, followed by post-fixation in 4% PFA/PBSTw for 20 min at RT and washed 5 x 5 min in PBSTw. They were forwarded to the probe detection stage and pre-hybridized with pre-warmed hybridization buffer for 30 min at 37 °C, then incubated overnight (>12 h) at 37 °C in probe solution containing 2 pmol of each probe set and 500 µL of pre-warmed hybridization buffer. Next, they were washed 2 x 5 min in 5x SSCT at RT to remove excess probes, and then pre-amplified with 500 µL of equilibrated amplification buffer for 30 min at RT. Meanwhile, 30 pmol of hairpin h1 and h2 were prepared separately by snap-cooling 2 µL of 3 µM stock at RT darkly for 30 min after 95 °C heat for 90 s. The pre-amplification solution was removed and replaced with the hairpin solution, and the mixture was incubated overnight (>12 h) at 37 °C. Excess hairpins were removed on the following date by washing in 5x SSCT for 4 x 20 min. The samples were stored long-term at 4 °C in dark condition until imaging.

#### RNA isolation and Quantitative reverse transcription PCR (qRT-PCR)

qPCR was performed on 4 dpf larvae from WT, *tenm4^+/-^*, *tenm4^-/-^* and *Tg(tenm4:Gal4*), with the primers targeting the downstream of KO and KI sites at *tenm4* exon 7 and exon 8. RNA extraction was conducted in Trizol reagent following the standard protocol. Fifteen larvae were pooled and homogenized in 200 μl of Trizol and 40 μl of chloroform using a pellet pestle. After vortexing and incubating for 2 min at RT, it was centrifuged at 12000 xg for 15 min at 4 °C to separate the phases, with the top aqueous phase transferred to a new tube. To precipitate RNA, it was mixed with 100 μl of 100% isopropanol plus 5 μg glycogen by inverting the tube 20 times, then incubated for 10 min at RT and centrifuged at 14000 xg for 10 min at 4 °C. The RNA pellet was washed twice with 1 ml of 75% ethanol by centrifuging at 7500 xg for 5 min at 4 °C, then air-dried at RT for 10 min and resuspended in 15 μl of DPEC-treated nuclease-free water. 1 μg RNA was reverse transcribed using SuperScript™ III First-Strand Synthesis System (Invitrogen) and 50 ng cDNA was applied to qPCR using FastStart Essential DNA Green Master (Roche), according to the supplier’s instructions. Relative RNA expression levels were calculated using the ddCt method compared to Gapdh and normalized to control samples. All primers were listed in Key Resource Table

#### Cryosectioning of retinal tissue

Cryosection experiment was performed on the retina from knock-in larvae *Tg(tenm4:Gal4;UAS:GFPCAAX*) at 5 dpf. Larvae were fixed and washed as described above. Their left and right retinae were dissected and placed into two separate tubes containing sucrose with an ascending concentration at 15% and 30% in PBS for 2 h each at RT, then 40% overnight at 4 °C with gentle shaking. Twelve cryo-protected retinae were transferred to the empty chamber, where the solution was removed and then was filled with OCT medium. The retinae were dorsally placed as a 4 x 3 array, and frozen on dry ice or at −80 °C in a sealed box. The cryo-sectioned retinae were cut to 20 μm using Leica CM1950 at −20 °C and stuck to Superfrost Plus slices, then long-term stored at −80 °C.

#### Immunohistochemistry

IHC experiments were performed on 5 dpf cryo-sectioned retinae using anti-GFP, as well as PFA-fixed whole-mount WT and *tenm4^-/-^* larvae from 3 dpf to 5 dpf using rabbit anti-PH3. Each sample was rehydrated/washed in PBS/0.8%Triton (PBST) for 3 x 10 min and then blocked in 3% NGS and 1% BSA in 0.8% PBST for 1h at RT. Primary antibodies were diluted to 1:1000 in blocking solution and incubated with retinae slices or larvae overnight at 4 °C. After washed with PBST for 6 x 20 min, retinae slices or larvae were incubated with secondary antibodies (Goat anti-chicken Alexa Fluor 488 or Goat anti-rabbit Alexa Fluor 546, Invitrogen) and DAPI in the blocking solution (1:1000) at RT for 3 h or overnight at 4 °C, respectively, followed by washed with PBST for 6 x 20 min. Three droplets of Fluoromount-G medium were added to the slice, which was then covered with a glass coverslip coated with nail polish to prevent dehydration. The slices and larvae (in PBS for short-term storage or glycerol for long-term storage) were stored at 4 °C in the dark until imaging.

#### Whole Mount TUNEL assay

TUNEL assay was performed on WT and *tenm4^-/-^* larvae from 3 dpf to 5 dpf according to the supplier’s instructions (Promega). Larvae were rinsed in PBS for 4 x 5 min and fixed in 4% PFA/PBST for 10 min at RT. After washing with PBST for 4 x 5 min, fixed larvae were equilibrated in equilibration buffer overnight at RT, followed by incubated in rTdT incubation buffer overnight at 4 °C. To stop the reaction, larvae were immersed in 2x SSC for 20 min at RT and washed in PBS for 2 x 5 min, then in PBST for 5 min. The samples were stored at 4 °C in dark conditions until imaging.

#### Sparse labelling of retinal ganglion cells

Sparse labelling of RGCs was conducted for time-lapse, structural, and glutamate imaging. The activator vector *ath5:Gal4* and the effector vector *UAS:Synaptophysin_GFP;UAS:DsRedx*, and pTol2-10xUAS:SFiGluSnFR;cmcl2:mCherry were prepared using QIAprep Spin Miniprep Kit (Qiagen). Each Gal4-UAS vector combination was co-injected at a concentration of 30 ng/µl in Danieau solution into WT and *tenm4^-/-^*embryos at 1-4 cell stage. Only larvae showing sparsely labelled RGCs expressing both Syp:GFP and DsRedx, or iGluSnFR in optic tectum were caught for analysis. For structural imaging, the positive larvae imaged at 4 dpf were cultured separately in 35 mm petri dishes and then imaged again at 10 dpf, while time-lapse and glutamate imaging was conducted only at 3 dpf or 4 dpf, respectively.

#### Confocal imaging

Live or fixed positive larvae were mounted dorsally using 1% LMP agarose in Danieau or PBS, respectively. Structural imaging of 6 dpf fixed whole brain, as well as 4 dpf and 10 dpf RGCs in the optic tectum was conducted using LSM 880 confocal microscope with a 20x/1.0 NA water-immersion objective (Zeiss), and with additional equipped incubator setting at 28.5 °C for time-lapse imaging of RGCs over 10 hours with 10 min intervals from 3 dpf to 3.5 dpf. Other fixed larvae and retina were conducted using LSM 800 confocal microscope with a 40x/1.3 NA oil-immersion objective. Laser was excited by 405 nm (DAPI), 488 nm (GFP), 568 nm (*gad1b* probe and DsRedx), 633 nm (*tenm4* probe). Optical sections were obtained at 0.66 µm or 1 µm intervals through the z-axis and 1 AU pinhole aperture. Tiled z-stacks were acquired with 10% overlap for whole-brain imaging and stitched using Zen Black. Maximum intensity projections and 3D rotated images were generated using ImageJ.^90^

#### Image registration

Registration of images from HCR and transgenic lines was performed using the ANTs toolbox version 2.6.4.^89^ Images were converted to Nrrd format using ImageJ. The HCR images of *gad1b* were used as the template and registered to align to the mapzebrain coordinate system^35,36^ using the reference brain fixed_standard_GAD1b.nrrd with the following code: antsRegistration -d 3 --float 1 -o [fish1_, fish1_Warped.nii.gz] -n WelchWindowedSinc --use-histogram-matching 0 -r [fixed_standard_GAD1b.nrrd, fish1_Gad1b.nrrd,1] -t Rigid[0.1] -m MI[fixed_standard_GAD1b.nrrd, fish1_Gad1b.nrrd,1,32,Regular,0.25] -c [200x200x200x0,1e- 8,10] -f 12x8x4x2 -s 4x3x2x1vox -t Affine[0.1] -m MI[fixed_standard_GAD1b.nrrd, fish1_Gad1b.nrrd,1,32,Regular,0.25] -c [200x200x200x0,1e-8,10] -f 12x8x4x2 -s 4x3x2x1vox -t SyN[0.05,6,0.5] -m CC[fixed_standard_GAD1b.nrrd, fish1_Gad1b.nrrd,1,2] -c [200x200x200x200x10,1e-7,10] -f 12x8x4x2x1 -s 4x3x2x1x0

The deformation matrices computed above were then applied to any other image channel using the following ANTs command: antsApplyTransforms -d 3 --float 1 -n WelchWindowedSinc -i fish1_HCR.nrrd -r fixed_standard_GAD1b.nrrd -o fish1_HCR_Warped.nii -t fish1_1Warp.nii.gz -t fish1_0GenericAffine.mat The above registration was repeated for another two fish. The three images were then arithmetically averaged using ImageJ,^90^ which are uploaded to mapzebrain.

#### In vivo calcium Imaging

Measurement of visual voxel responses indicated by SyGCaMP3 (Ca^2+^ releasing) on RGCs was performed on Tg(*Isl2b:Gal4; UAS:SyGCaMP3; mitfa*) in WT and *tenm4^-/-^* background at 4 dpf, following the established protocol.^50^ Non-anaesthetised larvae were dorsally immobilized in 2% LMP agarose (prepared in Danieau) on the edge of a raised glass platform with a single eye out of agarose, allowing free-moving and unobstructed view to the stimuli. After solidifying, the mounted larvae were placed in a custom-made dark box filled with Danieau for over 30 min to allow for acclimation, minimizing larval drift and movement. Recordings were completed in the afternoon.

Confocal functional Imaging were performed using an LSM 710 microscope with a spectral detection scan head and 20x/1.0 NA water-immersion objective (Zeiss), and a DLP Pico Projector (Optoma). Lavae was placed in the chamber filled with Danieau to record the response of RGC projections in 2-4 Z-planes around the middle plane of optic tectum, separated by 2 μm. Each recording was repeated twice. Functional time-series visually evoked calcium responses consisted of 1200 frames at a rate of 4.1 Hz, at a resolution of 0.415×0.415 µm resolution (256× 256 pixels), at an excitation wavelength of 488 nm, and a pinhole aperture of 1AU. The stimuli of moving gratings were created through a custom written LabView and MATLAB (MathWorks) code running through a ViSaGe stimulus presenter (Cambridge Research Systems) and projected laterally onto a diffusion filter (3026, Rosco) bonded to one side of the chamber. The grating was comprised of 12 different orientations of black and white bar (100% contrast) at 30° intervals, with each randomly presented for 3 seconds with a 10 second break with a blank white screen to enable signals to return to baseline.

#### In vivo 2-photon glutamate imaging

Glutamate release from RGCs was measured at single cell level in *tenm4^+/+^* and *tenm4^-/-^* larvae at 4dpf. Non-anaesthetised larvae were dorsally immobilized in 2% LMP agarose (prepared in Danieau) on top of a glass coverslip, which then placed in a black box filled with Danieau. Eye movements were further prevented by injection of α-bungarotoxin (1 nL of 2 mg/ml) into the ocular muscles behind the eye.

Two-photon functional imaging was performed in the afternoon using an A1R MP microscope equipped with a 4-channel GaAsP NDD and a 25×/1.1 NA water-immersion objective (Nikon). Visual stimuli consist of moving gratings (100% contrast of black and white bar) were generated and controlled using PsychoPy and delivered through an LCD screen poisoned underneath a custom-made Perspex chamber. A long-pass red glass filter (FGL610, Thorlabs) was poisoned between the LCD screen and the chamber to allow for simultaneous imaging and visual stimulation. Each stimulus was presented for 11 s. Functional time-series of visually evoked glutamate responses consisted of ∼2000 frames at a rate of 7.8 Hz, at a resolution of 0.397×0.397μm resolution (256×128 pixels), at an excitation wavelength of 930 nm provided by a Chameleon Ultra II Mode-locked titanium-sapphire laser (Coherent).

#### Optokinetic response test

OKR test was collaborated with Isaac Bianco’s Lab (UCL, UK).^92^ 5 dpf old larvae derived from incrossed *tenm4^+/-^*mutants were embedded in 3.5% LMP agarose with eye free-moving and immersed in filtered fish facility water in 35 mm peri dishes. They were then placed in the culturing incubator for over 2 hours to recover from anesthesia and for acclimatization. In the OKR test, 12 unique rotating stimuli, varying in spatial frequencies (12, 24, 36, 48, 60), angular velocities (10, 30, 60, 90 °/s), and contrast (0.05, 0.1, 0.25, 0.5, 1), were controlled by LabVIEW (National Instruments) and orderly presented in the drum covered by diffusion paper around the embedded larvae. Each stimulus was repeated twice. AVT Pike cameras were used to locate larvae by NImax and track horizontal eye movement (100 frames per second) under infrared illumination (850 nm), which was recorded by LabVIEW. All assays were done blind to genotype with post-experimental genotyping.

#### Optomotor response test

OMR test in free swimming larvae was modified from the established closed-loop paradigm.^54^ Individual 5 dpf old larvae derived from incrossed *tenm4^+/-^* mutants were placed in 120 mm circular dishes containing filtered fish facility water and acclimated for 15 min. The stimuli, varying in stripe width (0.01, 0.02, 0.04, 0.08, 0.16 AU) and contrast (0.4, 1), were controlled by custom-made software in Python 3.11 and OpenCV 4.1 and were projected in an orderly manner onto a transparent, infrared-illuminated (940 nm) dish base covered with diffusion paper. Each stimulus was repeated three times, with 25 seconds of motion at 0.05 cm/s and 5 seconds of rest, in either the leftward or rightward direction perpendicular to fish’s orientation. Larvae were tracked by high-speed CMOS camera (Basler acA2040-90 μm-NIR) with a zoom lens (Zoom 7000, 18-108 mm) at 90 fps, and their position and orientation were recorded based on the overall body shape, that is the difference between each current frame and the average over the frames corresponding to the last 800s and updated every 50 frames through background subtraction. Bouts were defined as turns if the absolute orientation change exceeded 2°. All assays were done blind to genotype with post-experimental genotyping.

### QUANTIFICATION AND STATISTICAL ANALYSIS

Descriptive statistics including normality testing were completed to determine appropriate statistical methods. All statistical tests were two-tailed and conducted using GraphPad Prism 10 or RStudio 4.4.3, with the type and N described in the figure legends. P-value for multiple comparisons was corrected by Geisser-Greenhouse unless otherwise noted, and significance was set to p<0.05*, p<0.01**, p<0.001***, p<0.0001****. All precision measurements are summarized in Table S1.

#### Cell distributional and fluorescent intensity analysis

Cell numbers were counted using cell counter in ImageJ. The percentage of RGCs was measured and averaged by counting *tenm4*^+^ RGCs and DAPI stained nuclei in one superficial section and deep section of 1µm. The number of TCs were counted in a superficial stack and deep stack (20μm each), separated by an interval of 30μm.

The fluorescent intensity profile of retina and optic tectum of *tenm4* HCR and reporter line was plotted using ImageJ through “Plot Profile” function on a square. The regions including each AFs of RGC downloaded from mapzebrain were outlined using “Binary - Outline” function in ImageJ, with the intensity measured through “Analyze Particles” function.

#### Morphological analysis

Morphology of RGCs from sparse labelling was analyzed blindly using Imaris 9.1 (Bitplane) through surface, spot, filament with Sholl analysis function. The entire 3D image was imported as maximum intensity projections. The region of interest (ROI) was selected and masked based on the cell arbor channel using ‘surface’ module, in which the intensity of channels was measured. Since they are determined by the amount of vector expressed in each cell and two channels are independent, synaptic puncta fluorescence was normalized to cell arbor signal using the following formula through channel arithmetic default plugin: *_Normalized Intensity_*=*_Matrix_* (*_red_* or *_green c_*ℎ*_annel_*)/*_Max intensity_* (*_red c_*ℎ*_annel_*). Next, another ‘surface’ module was applied to new masked channel to calculate the volume of each punctum in RGCs while their number was quantified using ‘spots’ module. All steps were quantified automatically using the same settings to ensure an unbiased measurement. The length of RGC branches were calculate using ‘filament’ module with the first branching point as the start point, then Sholl analysis was applied based on the reconstructed axonal arbor.

For time-lapse imaging, puncta number and branch length at each 10 min for a short 3 h period 3 h and each 1 h for long 10 h period were measured, thereby puncta turnover and branch motility, indicated by the absolute difference of puncta density or branch length between two consecutive time points, were calculated and normalized to the number of stable branches that quantified as follows. Axonogenesis was analyzed using 4D SPA.^47^ The full filament data was exported and converted hoc to bcf file in Command Prompt, then imported to 4D SPA. Each branch at consecutive time points was manually designated as either ‘matched’ or ‘retracted’ according to the software’s suggestion. Once the alignment was completed for the pair, the output results were acquired to perform MultiplePairAnalysis in Command Prompt, which summaries the whole branching events, categorized as retracted (present at one time-point but lost at the next), transient (lost later), and newly-added branches (appearing after the first time point and persisting to the endpoint). They were also normalized to the number of stable branches. Additionally, newly generated branches in each time interval were indicated in distinct color, and the fraction of intermittent branches (previously classified as stable branches but final retracted) was quantified by these color-coded branches.

#### Functional analysis

Functional calcium time-series images were analyzed based on previous script.^50^ The images were preprocessed using the SPM12 through rigid-body algorithm for motion correction as follows. Median filtered with a kernel size of 1 voxel (0.415 μm) and spatially smoothed with the Gaussian smoothing kernel of 2 voxels were used to minimize background noise (dark and shot) and improve the signal-to-noise ratio, respectively. Next, low-frequency baseline drifts in baseline (B) were corrected by cubic-spline algorithm interpolation computed from the inter-epoch 5s intervals. After preprocessing to create anatomical reference images, the relative signal intensity changes (ΔF = F-B) and the integral threshold (5×SDs) were calculated at each voxel for the population-level voxel wise analysis, with the latter determined from the variance of ΔF changes during the inter-epoch intervals and null condition. Total visually responsive was quantified from all voxels exceeding the empirical threshold. It was then categorized into direction-selective (DS) or orientation-selective (OS) subtypes by applying specific thresholds: OS = DSI < 0.5 and OSI > 0.5; DS = DSI > 0.5 and OSI < 0.5; with R^2^ > 0.8. DSI is calculated by comparing responses between the preferred direction (R_pref_) and its 180° opposite, or null direction (R_null_), defined by 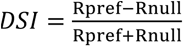. Similarly, OSI is derived from the response to the preferred orientation (R_pref_) relative to the orthogonal axis (R_orth_; 90° angular distance), defined by 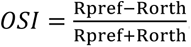

Functional glutamate release from RGCs was analyzed using Igor Pro 7.0 (WaveMetrics).^93^ Image sequences were preprocessed by background subtraction and frame alignment to correct for motion. Single-cell ROIs were detected automatically based on fluorescence variance and pixel correlation, followed by manual corrected to select ROI ROIs exhibiting a response exceeding predefined threshold of ΔF/F. To reduce the effect of photobleaching, an exponential decay function was fitted to the fluorescence signals and subtracted from the raw traces. 300 frames (∼38 s) of each recording were included in the analysis. Baseline fluorescence (F0) was defined as the mean signal during the 5s prior to stimulation. Responses were calculated as ΔF/F=(F-F0)/F0. For group analysis, the following metrics were extracted: peak amplitude: the maximum ΔF/F value; response latency: time from stimulus onset to the peak response. They were quantified between 9.5 s and 13 s (frames 75-103) after recording onset, corresponding to the early stimulus-evoked glutamate peak. The other metrics were analyzed between 9.5 s and 22 s (frames 75-173) encompassing the full stimulus-evoked glutamate transient: response area (area under curve): the trapezoidal integral of the ΔF/F signals (>0); decay time: the time (in seconds) from the peak response to return to 10% above baseline, computed from the post-peak portion of the signal; response frequency: total number of glutamate release events (peaks); response latency: time to trigger the peak response.

#### Behavioral analysis

For the OKR experiment, each eye movement trace was analyzed using a customized MATLAB script by automatically marking 1 s windows below a certain threshold. The sections of unstable saccades from the disruption of spontaneous horizontal saccades or jaw movements were excluded manually.^92^ The slow-phase gain (the ratio of eye velocity to stimulus velocity), saccadic eye velocity, and oculomotor range of left and right eye in each direction, were then computed.

The raw data from OMR experiment were pre-processed using Pycharm software to exclude first 10 seconds of data, allowing the fish for the acclimatization to the stimulus changes.^94^ Additionally, the tracking errors and aberrant behaviours (either hypoactivity or hyperactivity) were dropped, including large contour area (>2000 pixel), long interbout interval (>10s), fate swim speed (>6 cm/s), large angle for the absolute orientation changes (>150°), position closed to the wall (<0.5 cm). Any complete trials in which these cases exceeded 5% were eliminated from the analysis. The bout speed, duration, distance, number, interbout interval and fraction of correct turns, were then computed separately under vertical left or right stimuli at different contrast and stripe widths.

## Notes

### Competing Interest Statement

The authors have declared no competing interest.

