## Supplemental Figures and Tables for "Lateralized Tenm4 expression biases visual performance and modulates axonal growth and pre-synapse size in retinal ganglion cells"

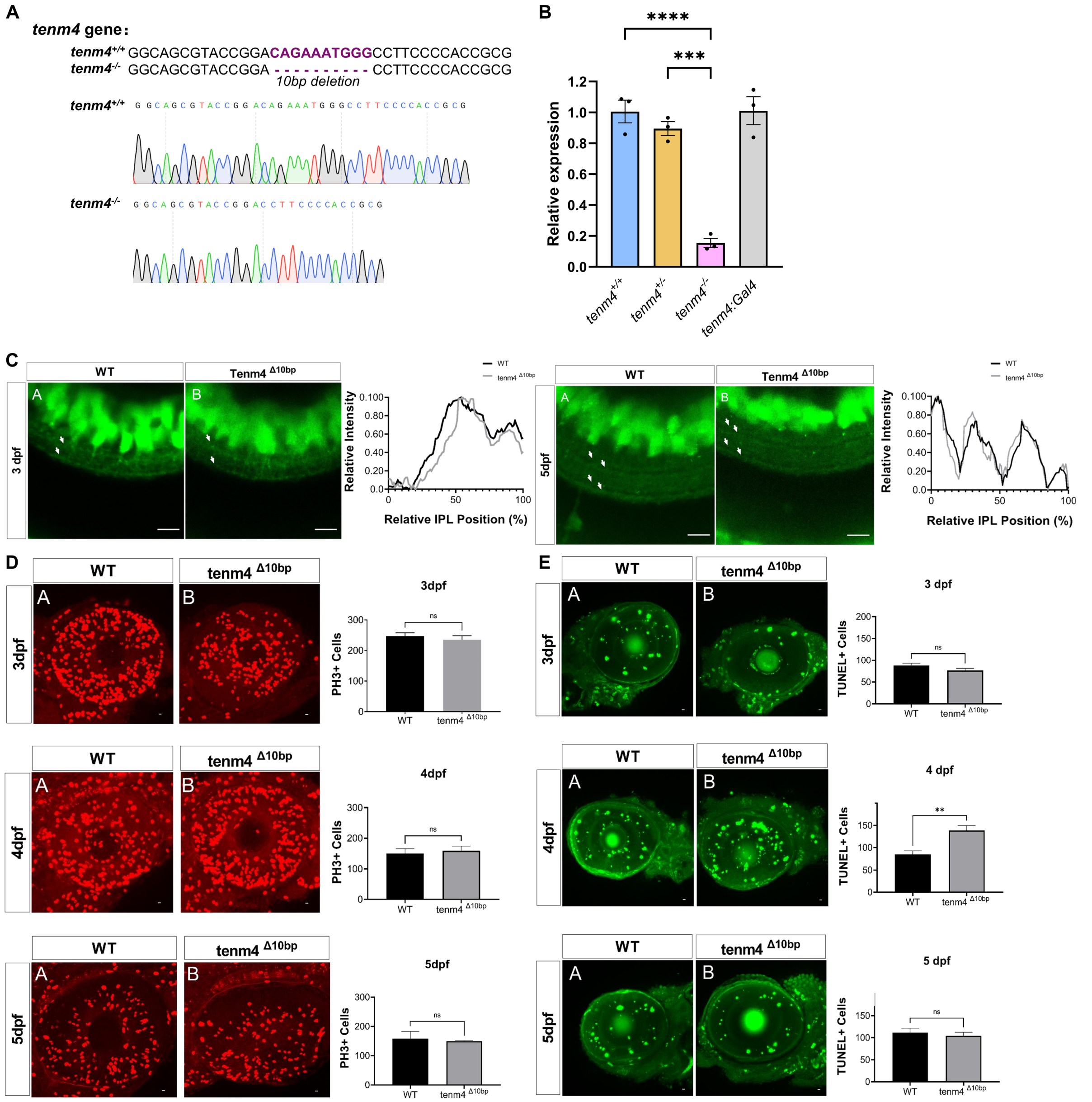


**Figure S1 related to Figure 1 and 2 in main text.**
**(A)** CRISPR-mediated genetic KO of *tenm4*. Sanger sequencing chromatograms show the 10bp deletion present in *tenm4*$^{-/-}$ mutant larvae. **(B)** Diagram shows the relative *tenm4* mRNA expression levels in *tenm4*$^{+/+}$, *tenm4*$^{+/-}$, *tenm4*$^{-/-}$ and *Tg(tenm4:Gal4)* animals. P-value is from unpaired t-test comparing *tenm4*$^{+/+}$ to the other genotypes. N= 3 biological and technical replicates for each genotype. **(C)** Inner plexiform layer (IPL) stratification pattern of RGCs the retinae of *tenm4*$^{+/+}$ and *tenm4*$^{-/-}$ animals crossed to the *Tg(Isl2b-Gal4;UAS-GFP)* line at 3 dpf and 5 dpf, with their relative intensity in IPL position. 0% corresponds to the inner nuclear layer (INL)/IPL boundary, whereas 100% corresponds to the IPL/GCL (ganglion cell layer) boundary. Comparison is done using two-way ANOVA. **(D and E)** Confocal images and comparison of PH3-stained mitotic cells (red, D) and TUNEL stained apoptotic cells (green, E) in *tenm4*$^{+/+}$ and *tenm4*$^{-/-}$ retina at 3, 4 and 5 dpf. Scale bar: 10μm. n=5 for each genotype. P-values are from unpaired t-test.

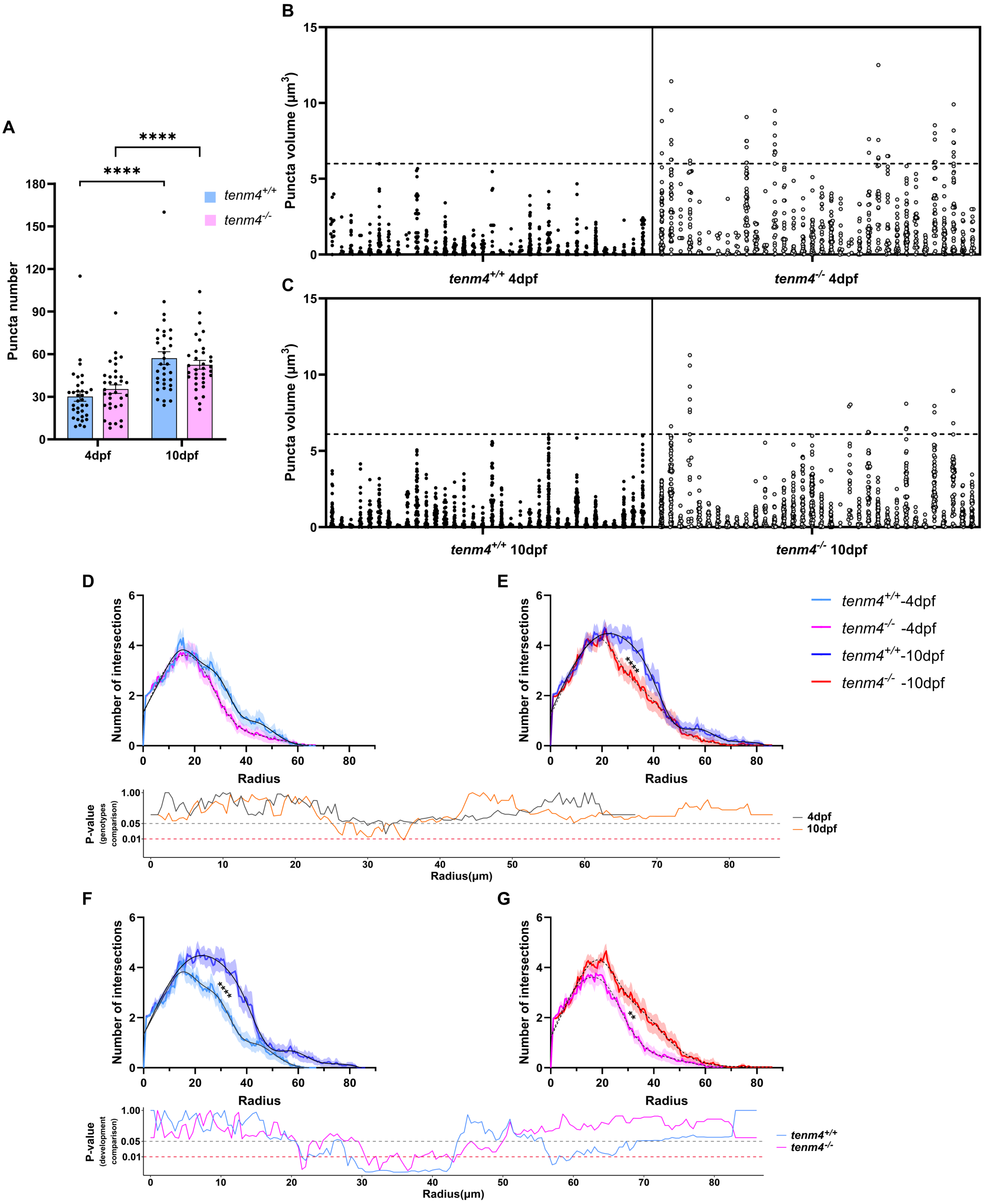


**Figure S2 related to Figure 2 in the main text.**

**(A)** Quantification of presynaptic puncta number in *tenm4*$^{+/+}$ and *tenm4*$^{-/-}$ RGCs at 4 dpf and 10 dpf. Each dot represents values from one individual, n= 34 for each genotype; p-value is from two-way ANOVA repeated comparison without correction and Fisher's LSD post-hoc test. All error bars are mean ± S.E.M. **(B and C)** Individual punctum volumes for each *tenm4*$^{+/+}$ (left) and *tenm4*$^{-/-}$ (right) RGC at 4 dpf (B) and 10 dpf (C). Dashed line indicates the maximal presynaptic volume found in wildtype animals (*tenm4*$^{+/+}$). **(D to G)** Sholl analysis of *tenm4*$^{+/+}$ and *tenm4*$^{-/-}$ RGC arborizations. Comparisons are done across genotypes at 4 dpf (D) and 10 dpf (E) with p-value below, or within genotypes during development for *tenm4*$^{+/+}$ (F) and *tenm4*$^{-/-}$ (G) RGCs, with p-values indicated below. The black curve indicates *tenm4*$^{+/+}$ vs *tenm4*$^{-/-}$ at 4 dpf; the orange curve indicates *tenm4*$^{+/+}$ vs *tenm4*$^{-/-}$ at 10 dpf; the light blue curve indicates *tenm4*$^{+/+}$ RGC at 4 dpf versus 10 dpf; the magenta curve indicates *tenm4*$^{-/-}$ RGC at 4 dpf versus 10 dpf. P-values are calculated using three-way ANOVA repeated comparison with Tukey's post-hoc test.

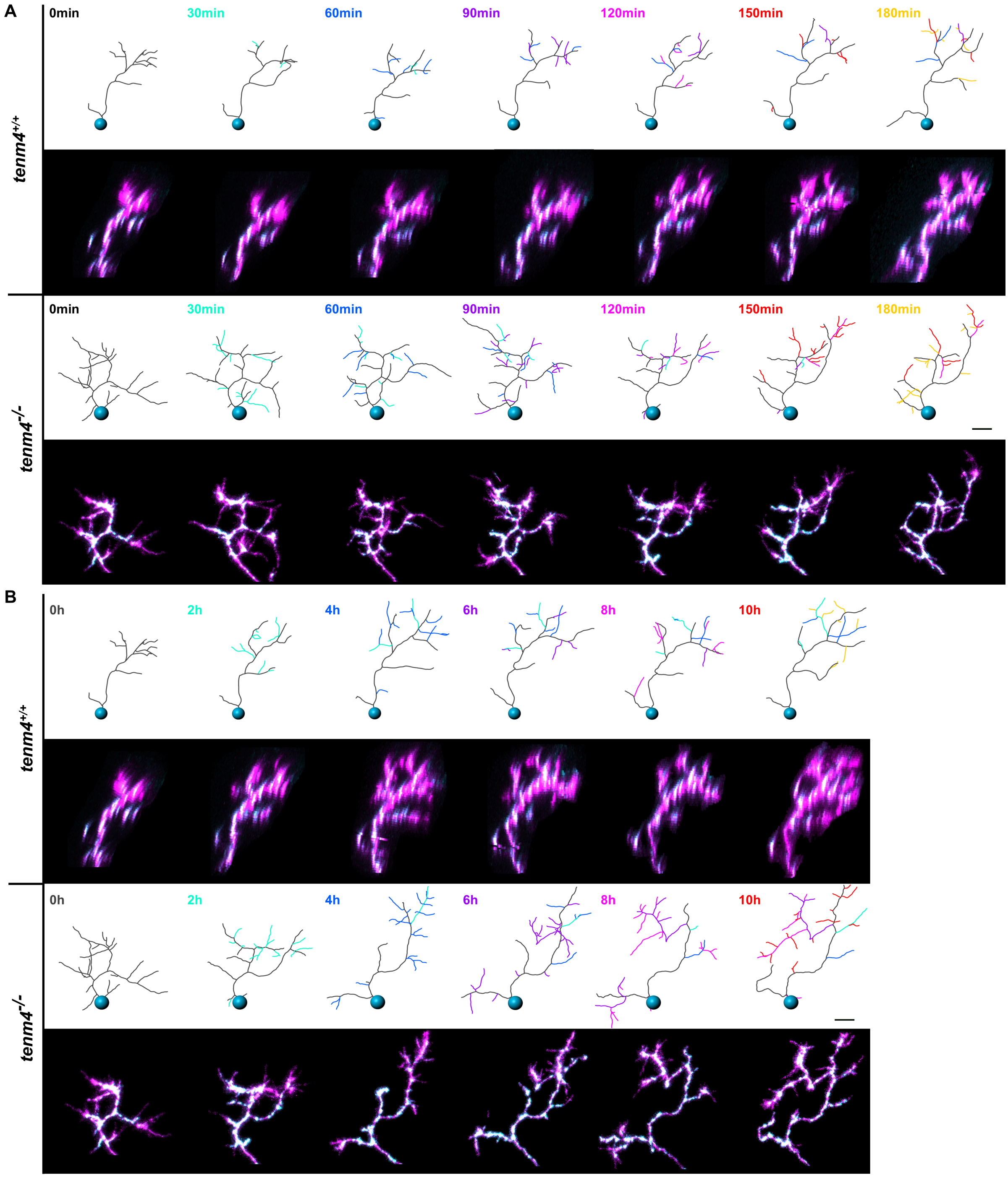


**Figure S3 related to Figure 3 in the main text and Movie S1 and S2.**

Overview of confocal time-lapse images of *tenm4*$^{+/+}$ and *tenm4*$^{-/-}$ RGC from 3 to 3.5 dpf in the initial 3h window **(A)** and in the 10 h window **(B)**, with their 3D-reconstructions shown above of each confocal image. Cell arbors are shown in magenta, while synaptophysin puncta are shown in cyan. Newly generated branches are color-coded for each 30-minute window for the first 3 hours and then for 2 hour windows until the end of the 10 hour time-lapse. Scale bar: 10μm.

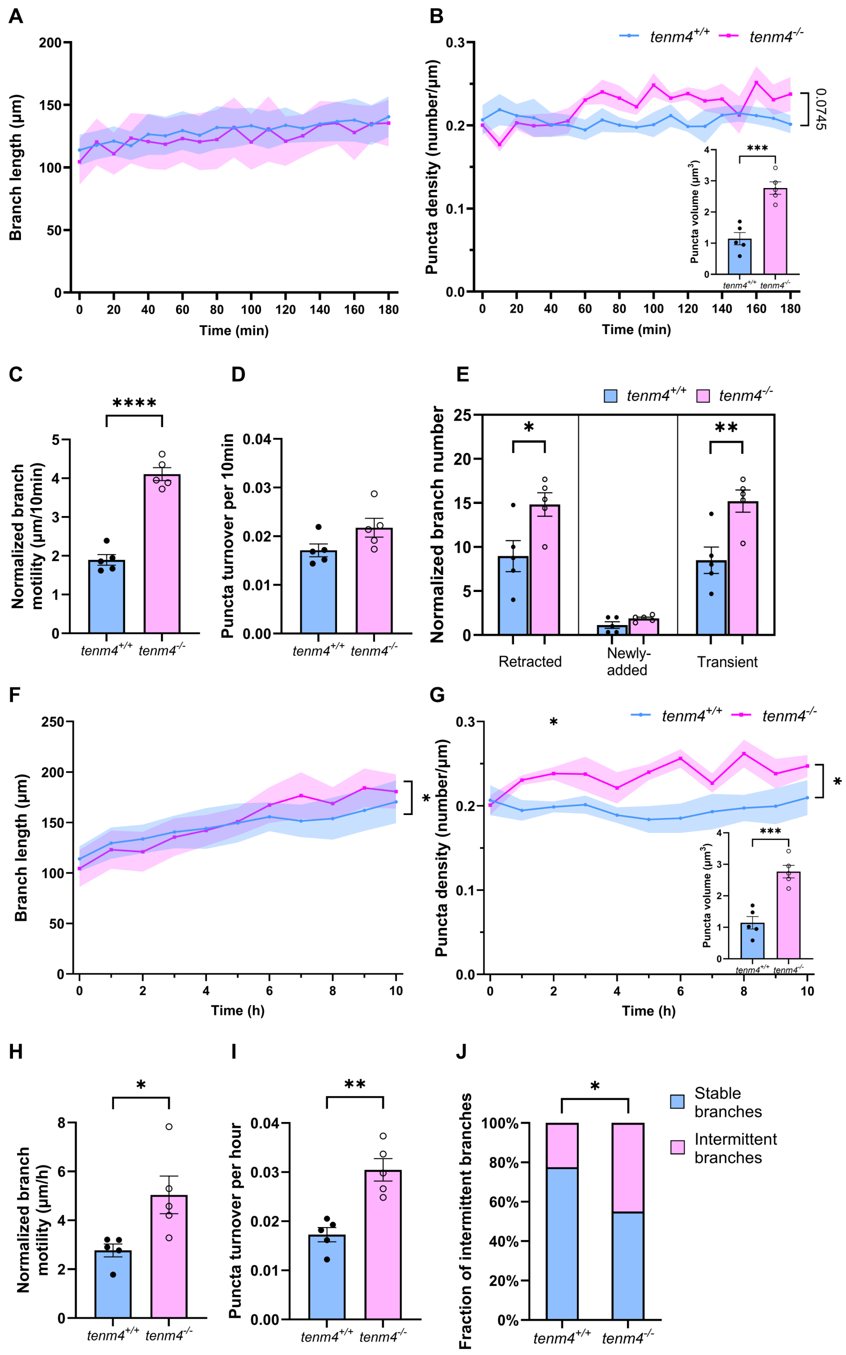


**Figure S4 related to Figure 3 in the main text.**

*Tenm4*$^{-/-}$ RGCs with enlarged puncta are selected to compare axonogenesis and synaptogenesis to *tenm4*$^{+/+}$ RGCs between 3 and 3.5 dpf. The growth curve of axonal branch and presynaptic puncta density, as well as normalized branch motility and puncta turnover, indicated by the absolute difference of branch length or puncta density between two consecutive time points (normalized to the number of stable branches), are compared in the first 3 h with 10 min intervals **(A to D)** and 10 h with 1 h intervals **(B to I)**. The number of branching events including retracted (present at one time-point but lost at the next), transient (lost later), and newly-added branches (appearing after the first time-point and persisting to the endpoint) are normalized to the number of stable branches (persisting over 4h) and compared in the first 3 h (F). The fraction of transient branches (initially classified as stable branches but eventually retracted) in 10 h, indicated by the number of transient branches over the number of stable branches, are compared in 10h (J). P-values for growth curves are from two-way ANOVA repeated comparison with Šidák post-hoc test, while others are from unpaired t-test. All error bars are mean ± S.E.M. n=5 for each genotype.

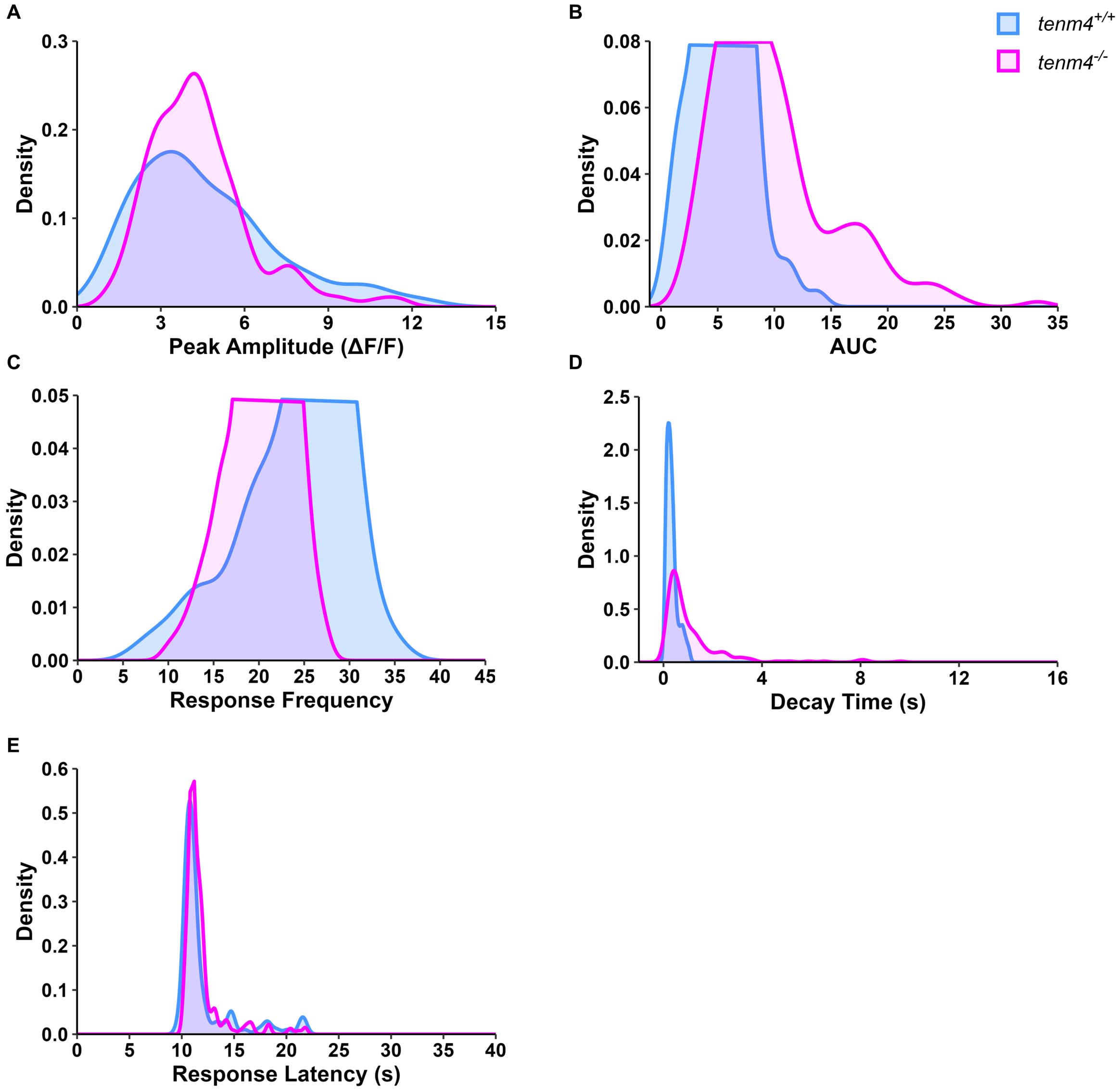


**Figure S5 related to Figure 4 in the main text.**
Distribution of relative frequency distribution of glutamatergic puncta (density) in *tenm4*$^{+/+}$ RGCs (blue) and *tenm4*$^{-/-}$ RGCs (magenta) for each response parameter: **(A)** peak amplitude: peak response of change in fluorescence ($\Delta F/F_0$); **(B)** area under the curve (AUC): cumulative response of the ΔF/F signals (>0); **(C)** response frequency: total number of glutamate release events (peaks); **(D)** decay time: the duration from the peak response to return to 10% above baseline; **(E)** response latency: time from stimulus onset to the peak response.

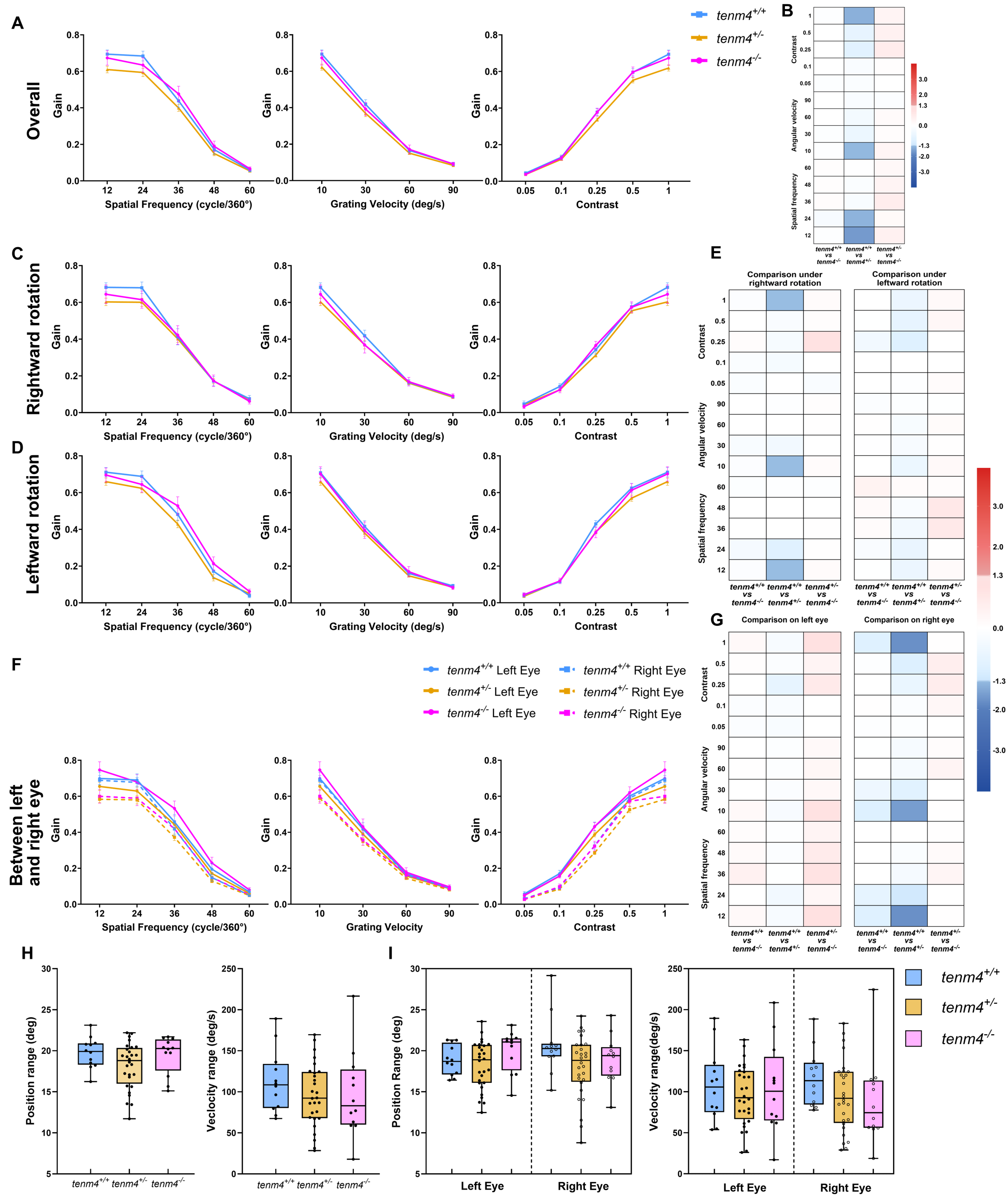


**Figure S6 related to Figure 5 in the main text.**

**(A to G)** The OKR gain, ratio of eye velocity to stimulus velocity, tested in 5 dpf larvae from incrossed *tenm4*$^{+/-}$animals under different stimuli, with changing spatial frequencies (12, 24, 36, 48, 60), angular velocities (10, 30, 60, 90 °/s), and contrast (0.05, 0.1, 0.25, 0.5, 1). The comparisons among three genotypes include overall comparison (A) with the heat map of p-values shown in (B), comparison under rightward rotation (C) and leftward rotation (D) with heat map of p-values shown in (E); comparison between the left and the right eye (F), with a heat map of p-values shown in (G). In the heat maps, the three columns depict *tenm4*$^{+/+}$ versus *tenm4*$^{-/-}$, *tenm4*$^{+/+}$ versus *tenm4*$^{+/-}$, *tenm4*$^{+/-}$ versus *tenm4*$^{-/-}$ comparisons. For heatmaps (B, E, G): A blue color indicates the performance of *tenm4*$^{+/+}$ is better than any of the mutants or the *tenm4*$^{+/-}$ is better than the *tenm4*$^{-/-}$. A red color indicates the opposite. p-values are from two-way ANOVA repeated comparison with Tukey post-hoc test. All error bars are mean ± S.E.M.

**(H and I)** The box chart showing the oculomotor ranges (min, max and mean) of overall position and velocity (H), as well as their separated comparison between left and right eye (I) in OKR test among three genotypes. Comparison is one-way repeated ANOVA with Šidák pot-hoc test. n=12 for *tenm4*$^{+/+}$ and *tenm4*$^{-/-}$, n=28 for *tenm4*$^{+/-}$.

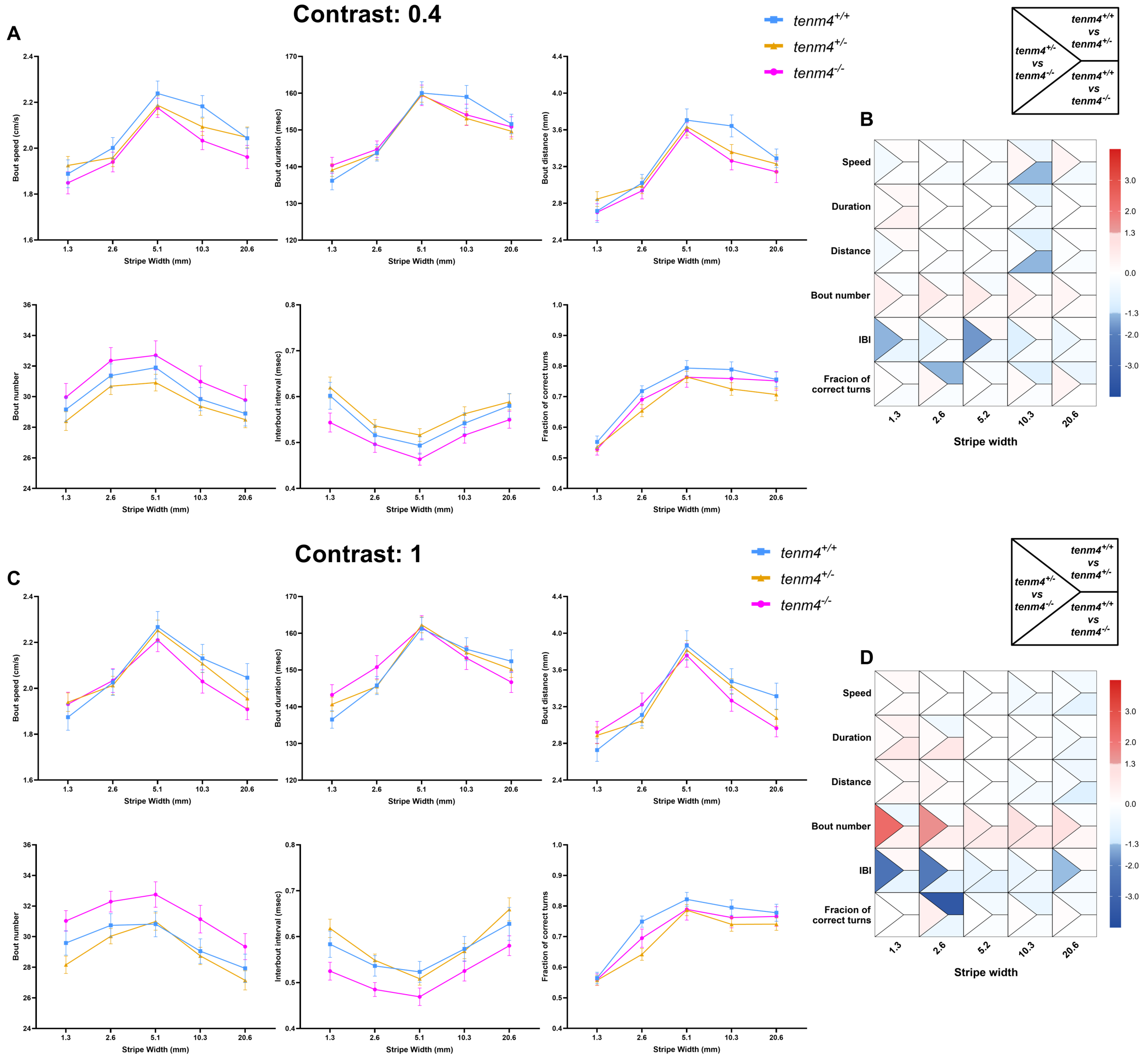


**Figure S7 related to Figure 5 in the main text.**

The overall fish OMR behavior, including bout speed, duration, distance, number, interbout interval and fraction of correct turns, testing in 5dpf larvae from *tenm4*$^{+/-}$ incross under grating varying in stripe width (0.01, 0.02, 0.04, 0.08, 0.16 AU) and contrast at 0.4 **(A)** and 1 **(C)**, with right heat map of p-value **(C and D**, respectively). Each cell is divided into a left triangle (*tenm4*$^{+/+}$ versus *tenm4*$^{-/-}$), an upper right trapezoid (*tenm4*$^{+/+}$ versus *tenm4*$^{+/-}$) and a bottom left trapezoid (*tenm4*$^{+/-}$ versus *tenm4*$^{-/-}$). For heatmaps (B, D): A blue color indicates the performance of *tenm4*$^{+/+}$ is better than any of the mutants (trapezoids) or the *tenm4*$^{+/-}$ is better than the *tenm4*$^{-/-}$ (triangle). A red color indicates the opposite. p-values are from two-way ANOVA repeated comparison with Tukey post-hoc test. All error bars are mean ± S.E.M. n=34 for *tenm4*$^{+/+}$, n=28 for *tenm4*$^{-/-}$, n=54 for *tenm4*$^{+/-}$.

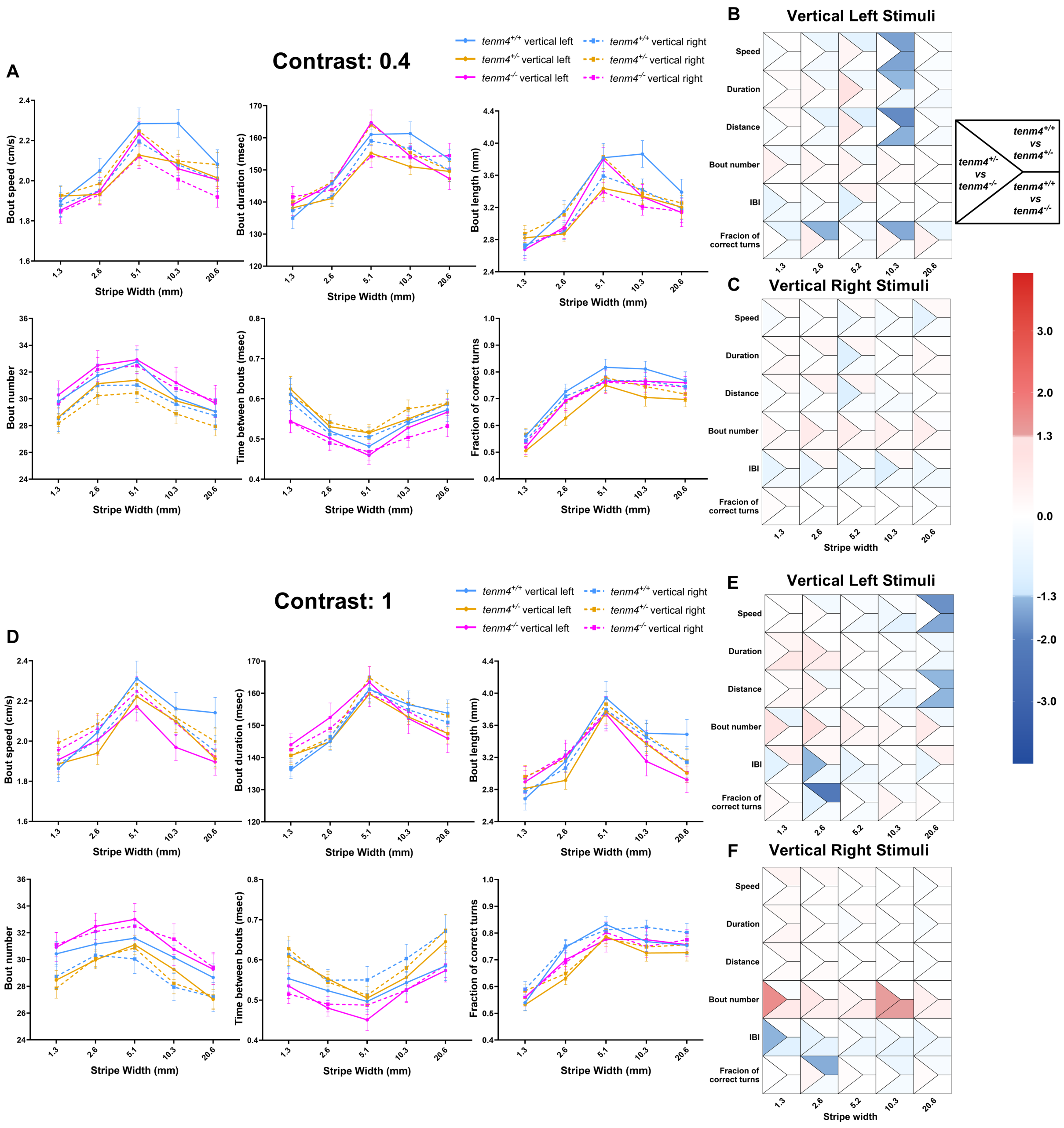


**Figure S8 related to Figure 5 in the main text.**

The separated fish OMR behavior, including bout speed, duration, distance, number, interbout interval and fraction of correct turns, under vertical left and right stimuli. The 5 dpf-tested larvae are derived from incrossing *tenm4*$^{+/-}$ animals, presented with gratings varying in stripe width (0.01, 0.02, 0.04, 0.08, 0.16 AU) and contrast at 0.4 **(A)** and 1 **(D)**. The heat map of p-values is to the right (**B, C** and **E, F**, respectively). Each area is divided into a left triangle (*tenm4*$^{+/+}$ versus *tenm4*$^{-/-}$), an upper right trapezoid (*tenm4*$^{+/+}$ versus *tenm4*$^{+/-}$) and a bottom left trapezoid (*tenm4*$^{+/-}$ versus *tenm4*$^{-/-}$). For heatmaps (B, C, E, F): A blue color indicates the performance of *tenm4*$^{+/+}$ is better than any of the mutants (trapezoids) or the *tenm4*$^{+/-}$ is better than the *tenm4*$^{-/-}$ (triangle). A red color indicates the opposite. p-values are from two-way ANOVA repeated comparison with Tukey post-hoc test. All error bars are mean ± S.E.M. n=34 for *tenm4*$^{+/+}$, n=28 for *tenm4*$^{-/-}$, n=54 for *tenm4*$^{+/-}$.

**Table S1, related to Figure 1-4 in the main text and Figure S1 : showing mean ± SEM**

| ***Figure*** | ***Description*** | ***mean ± SEM*** |
|---|---|---|
| Figure 1H | Left retina – dorsal part | 28.40% ± 2.10% |
| Figure 1H | Right retina – dorsal part | 32.87% ± 2.00% |
| Figure 1H | Left retina – ventral part | 30.83% ± 2.53% |
| Figure 1H | Right retina – ventral part | 37.55% ± 2.36% |
| Figure 1H | Left retina – nasal part | 30.69% ± 2.09% |
| Figure 1H | Right retina – nasal part | 36.37% ± 1.85% |
| Figure 1H | Left retina – temporal part | 28.05% ± 2.49% |
| Figure 1H | Right retina – temporal part | 34.59% ± 2.85% |
| Figure 1H | Left retina - overall | 29.30% ± 1.99% |
| Figure 1H | Right retina – overall | 35.36% ± 2.03% |
| Figure 1G | Left tectum – deep layer | 178.9 ± 6.54 |
| Figure 1G | Right tectum – deep layer | 190.7 ± 5.52 |
| Figure 1G | Left tectum – superficial layer | 129.8 ± 5.64 |
| Figure 1G | Right tectum – superficial layer | 142.3 ± 5.80 |
| Figure 1G | Left tectum - overall | 308.7 ± 11.12 |
| Figure 1G | Right tectum – overall | 333.0 ± 8.73 |
| Figure 2D | *tenm4*$^{+/+}$ - 4dpf | 0.69 ± 0.09 |
| Figure 2D | *tenm4*$^{-/-}$ - 4dpf | 1.64 ± 0.17 |
| Figure 2D | *tenm4*$^{+/+}$ - 10dpf | 0.77 ± 0.10 |
| Figure 2D | *tenm4*$^{-/-}$ - 10dpf | 1.17 ± 0.18 |
| Figure 2E | *tenm4*$^{+/+}$ - 4dpf | 0.17 ± 0.01 |
| Figure 2E | *tenm4*$^{-/-}$ - 4dpf | 0.22 ± 0.01 |
| Figure 2E | *tenm4*$^{+/+}$ - 10dpf | 0.23 ± 0.01 |
| Figure 2E | *tenm4*$^{-/-}$ - 10dpf | 0.24 ± 0.01 |
| Figure 2F | *tenm4*$^{+/+}$ - 4dpf | 176.22 ± 14.71 |
| Figure 2F | *tenm4*$^{-/-}$ - 4dpf | 157.44 ± 10.90 |
| Figure 2F | *tenm4*$^{+/+}$ - 10dpf | 263.97 ± 18.73 |
| Figure 2F | *tenm4*$^{-/-}$ - 10dpf | 222.67 ± 13.14 |
| Figure 3C | *tenm4*$^{+/+}$ | 1.145 ± 0.20 |
| Figure 3C | *tenm4*$^{-/-}$ | 2.247 ± 0.29 |
| Figure 3D | *tenm4*$^{+/+}$ | 1.89 ± 0.14 |
| Figure 3D | *tenm4*$^{-/-}$ | 3.82 ± 0.28 |
| Figure 3E | *tenm4*$^{+/+}$ | 0.017 ± 0.0013 |
| Figure 3E | *tenm4*$^{-/-}$ | 0.023 ± 0.0022 |
| Figure 3F | *tenm4*$^{+/+}$ - retracted | 8.97 ± 1.76 |
| Figure 3F | *tenm4*$^{-/-}$ - retracted | 13.61 ± 1.03 |
| Figure 3F | *tenm4*$^{+/+}$ - newly added | 1.13 ± 0.37 |
| Figure 3F | *tenm4*$^{-/-}$ - newly added | 1.819 ± 0.12 |
| Figure 3F | *tenm4*$^{+/+}$ - transient | 8.48 ± 1.50 |
| Figure 3F | *tenm4*$^{-/-}$ - transient | 13.23 ± 1.26 |
| Figure 3J | *tenm4*$^{+/+}$ | 2.77 ± 0.26 |
| Figure 3J | *tenm4*$^{-/-}$ | 4.42 ± 0.59 |
| Figure 3K | *tenm4*$^{+/+}$ | 0.017 ± 0.0015 |
| Figure 3K | *tenm4*$^{-/-}$ | 0.027 ± 0.0031 |
| Figure 3L | *tenm4*$^{+/+}$ | 24.00% ± 6.78% |
| Figure 3L | *tenm4*$^{-/-}$ | 43.77% ± 1.34% |
| Figure 4C | *tenm4*$^{+/+}$ | 4.618 ± 0.252 |
| Figure 4C | *tenm4*$^{-/-}$ | 4.388 ± 0.125 |
| Figure 4D | *tenm4*$^{+/+}$ | 5.420 ± 0.247 |
| Figure 4D | *tenm4*$^{-/-}$ | 9.735 ± 0.355 |
| Figure 4E | *tenm4*$^{+/+}$ | 24.347 ± 0.565 |
| Figure 4E | *tenm4*$^{-/-}$ | 20.441 ± 0.214 |
| Figure 4F | *tenm4*$^{+/+}$ | 0.335 ± 0.022 |
| Figure 4F | *tenm4*$^{-/-}$ | 1.276 ± 0.137 |
| Figure 4H | *tenm4*$^{+/+}$ - total | 3,724 ± 892.2 |
| Figure 4H | *tenm4*$^{-/-}$ - total | 9,014 ± 1,412 |

| | | |
|---|---|---|
| Figure 4H | *tenm4*$^{+/+}$ - non-tuned | 3,380 ± 819.1 |
| Figure 4H | *tenm4*$^{-/-}$- non-tuned | 7,960 ± 1,224 |
| Figure 4H | *tenm4*$^{+/+}$ - DSI | 127.4 ± 39.68 |
| Figure 4H | *tenm4*$^{-/-}$ - DSI | 468.1 ± 111.8 |
| Figure 4H | *tenm4*$^{+/+}$ - OSI | 216.7 ± 65.96 |
| Figure 4H | *tenm4*$^{-/-}$ - OSI | 586.3 ± 144.9 |

**Table S2, related to Figure 5 in the main text: showing mean ± SEM**

| ***Figure*** | ***Description*** | ***mean ± SEM*** | | | | |
|---|---|---|---|---|---|---|
| Figure 5B – 5F | *tenm4*$^{+/+}$ - Spatial frequency = 12 | 0.69 ± 0.023 | 0.68 ± 0.025 | 0.71 ± 0.025 | 0.70 ± 0.035 | 0.69 ± 0.026 |
| Figure 5B – 5F | *tenm4*$^{+/+}$ - Spatial frequency = 24 | 0.68 ± 0.027 | 0.68 ± 0.031 | 0.69 ± 0.030 | 0.69 ± 0.034 | 0.68 ± 0.027 |
| Figure 5B – 5F | *tenm4*$^{+/+}$ - Spatial frequency = 36 | 0.44 ± 0.035 | 0.41 ± 0.044 | 0.48 ± 0.034 | 0.46 ± 0.041 | 0.42 ± 0.035 |
| Figure 5B – 5F | *tenm4*$^{+/+}$ - Spatial frequency = 48 | 0.17 ± 0.025 | 0.17 ± 0.025 | 0.17 ± 0.034 | 0.19 ± 0.031 | 0.15 ± 0.020 |
| Figure 5B – 5F | *tenm4*$^{+/+}$ - Spatial frequency = 60 | 0.06 ± 0.009 | 0.08 ± 0.017 | 0.04 ± 0.011 | 0.07 ± 0.014 | 0.05 ± 0.007 |
| Figure 5B – 5F | *tenm4*$^{+/-}$ - Spatial frequency = 12 | 0.61 ± 0.018 | 0.60 ± 0.020 | 0.66 ± 0.020 | 0.65 ± 0.022 | 0.58 ± 0.019 |
| Figure 5B – 5F | *tenm4*$^{+/-}$ - Spatial frequency = 24 | 0.59 ± 0.022 | 0.60 ± 0.022 | 0.62 ± 0.025 | 0.63 ± 0.027 | 0.58 ± 0.023 |
| Figure 5B – 5F | *tenm4*$^{+/-}$ - Spatial frequency = 36 | 0.40 ± 0.018 | 0.40 ± 0.020 | 0.43 ± 0.021 | 0.44 ± 0.021 | 0.37 ± 0.018 |
| Figure 5B – 5F | *tenm4*$^{+/-}$ - Spatial frequency = 48 | 0.15 ± 0.012 | 0.17 ± 0.014 | 0.14 ± 0.018 | 0.17 ± 0.017 | 0.13 ± 0.009 |
| Figure 5B – 5F | *tenm4*$^{+/-}$ - Spatial frequency = 60 | 0.05 ± 0.007 | 0.07 ± 0.008 | 0.05 ± 0.010 | 0.06 ± 0.008 | 0.05 ± 0.006 |
| Figure 5B – 5F | *tenm4*$^{-/-}$ - Spatial frequency = 12 | 0.67 ± 0.04 | 0.64 ± 0.051 | 0.70 ± 0.038 | 0.75 ± 0.046 | 0.60 ± 0.038 |
| Figure 5B – 5F | *tenm4*$^{-/-}$ - Spatial frequency = 24 | 0.63 ± 0.04 | 0.62 ± 0.047 | 0.64 ± 0.042 | 0.68 ± 0.041 | 0.59 ± 0.041 |
| Figure 5B – 5F | *tenm4*$^{-/-}$ - Spatial frequency = 36 | 0.48 ± 0.04 | 0.42 ± 0.051 | 0.53 ± 0.049 | 0.53 ± 0.041 | 0.42 ± 0.046 |
| Figure 5B – 5F | *tenm4*$^{-/-}$ - Spatial frequency = 48 | 0.19 ± 0.03 | 0.17 ± 0.031 | 0.21 ± 0.036 | 0.23 ± 0.032 | 0.15 ± 0.027 |
| Figure 5B – 5F | *tenm4*$^{-/-}$ - Spatial frequency = 60 | 0.07 ± 0.01 | 0.06 ± 0.014 | 0.06 ± 0.013 | 0.08 ± 0.013 | 0.05 ± 0.009 |
| Figure 5B – 5F | *tenm4*$^{+/+}$ - Angular velocity = 10 | 0.69 ± 0.023 | 0.68 ± 0.025 | 0.71 ± 0.024 | 0.70 ± 0.035 | 0.69± 0.026 |
| Figure 5B – 5F | *tenm4*$^{+/+}$ - Angular velocity = 30 | 0.42 ± 0.026 | 0.42 ± 0.030 | 0.42 ± 0.031 | 0.42 ± 0.036 | 0.42 ± 0.020 |
| Figure 5B – 5F | *tenm4*$^{+/+}$ - Angular velocity = 60 | 0.17 ± 0.010 | 0.17 ± 0.015 | 0.16 ± 0.012 | 0.17 ± 0.013 | 0.16 ± 0.008 |
| Figure 5B – 5F | *tenm4*$^{+/+}$ - Angular velocity = 90 | 0.09 ± 0.008 | 0.09 ± 0.008 | 0.09 ± 0.010 | 0.09 ± 0.009 | 0.09 ± 0.008 |
| Figure 5B – 5F | *tenm4*$^{+/-}$ - Angular velocity = 10 | 0.62 ± 0.017 | 0.60 ± 0.020 | 0.66 ± 0.020 | 0.66 ± 0.022 | 0.59 ± 0.019 |
| Figure 5B – 5F | *tenm4*$^{+/-}$ - Angular velocity = 30 | 0.37 ± 0.018 | 0.37 ± 0.022 | 0.38 ± 0.017 | 0.39 ± 0.018 | 0.35 ± 0.022 |

| | | | | | | |
|---|---|---|---|---|---|---|
| Figure 5B – 5F | *tenm4*$^{+/-}$ - Angular velocity = 60 | 0.15 ± 0.007 | 0.16 ± 0.011 | 0.15 ± 0.007 | 0.16 ± 0.008 | 0.14 ± 0.007 |
| Figure 5B – 5F | *tenm4*$^{+/-}$ - Angular velocity = 90 | 0.08 ± 0.004 | 0.08 ± 0.005 | 0.09 ± 0.005 | 0.09 ± 0.004 | 0.08 ± 0.004 |
| Figure 5B – 5F | *tenm4*$^{-/-}$ - Angular velocity = 10 | 0.67 ± 0.040 | 0.64 ± 0.051 | 0.70 ± 0.039 | 0.75 ± 0.046 | 0.60 ± 0.038 |
| Figure 5B – 5F | *tenm4*$^{-/-}$ - Angular velocity = 30 | 0.39 ± 0.036 | 0.37 ± 0.044 | 0.40 ± 0.044 | 0.43 ± 0.045 | 0.36 ± 0.028 |
| Figure 5B – 5F | *tenm4*$^{-/-}$ - Angular velocity = 60 | 0.17 ± 0.024 | 0.17 ± 0.023 | 0.17 ± 0.028 | 0.18 ± 0.025 | 0.16 ± 0.023 |
| Figure 5B – 5F | *tenm4*$^{-/-}$ - Angular velocity = 90 | 0.09 ± 0.010 | 0.09 ± 0.012 | 0.08 ± 0.010 | 0.10 ± 0.012 | 0.09 ± 0.009 |
| Figure 5B – 5F | *tenm4*$^{+/+}$ - Contrast = 0.05 | 0.04 ± 0.008 | 0.05 ± 0.016 | 0.04 ± 0.011 | 0.06 ± 0.013 | 0.03 ± 0.008 |
| Figure 5B – 5F | *tenm4*$^{+/+}$ - Contrast = 0.1 | 0.13 ± 0.012 | 0.14 ± 0.015 | 0.12 ± 0.011 | 0.17 ± 0.019 | 0.09 ± 0.008 |
| Figure 5B – 5F | *tenm4*$^{+/+}$ - Contrast = 0.25 | 0.38 ± 0.018 | 0.34 ± 0.025 | 0.43 ± 0.018 | 0.43 ± 0.025 | 0.32 ± 0.017 |
| Figure 5B – 5F | *tenm4*$^{+/+}$ - Contrast = 0.5 | 0.60 ± 0.020 | 0.58 ± 0.020 | 0.62 ± 0.026 | 0.60 ± 0.026 | 0.59 ± 0.022 |
| Figure 5B – 5F | *tenm4*$^{+/+}$ - Contrast = 1 | 0.69 ± 0.023 | 0.68 ± 0.025 | 0.71 ± 0.025 | 0.70 ± 0.035 | 0.69 ± 0.0026 |
| Figure 5B – 5F | *tenm4*$^{+/-}$ - Contrast = 0.05 | 0.04 ± 0.003 | 0.04 ± 0.007 | 0.04 ± 0.006 | 0.05 ± 0.004 | 0.03 ± 0.003 |
| Figure 5B – 5F | *tenm4*$^{+/-}$ - Contrast = 0.1 | 0.12 ± 0.004 | 0.12 ± 0.006 | 0.12 ± 0.007 | 0.16 ± 0.006 | 0.09 ± 0.005 |
| Figure 5B – 5F | *tenm4*$^{+/-}$ - Contrast = 0.25 | 0.34 ± 0.009 | 0.31 ± 0.011 | 0.39 ± 0.010 | 0.39 ± 0.013 | 0.29 ± 0.01 |
| Figure 5B – 5F | *tenm4*$^{+/-}$ - Contrast = 0.5 | 0.55 ± 0.014 | 0.55 ± 0.014 | 0.57 ± 0.017 | 0.58 ± 0.018 | 0.53 ± 0.017 |
| Figure 5B – 5F | *tenm4*$^{+/-}$ - Contrast = 1 | 0.62 ± 0.017 | 0.60 ± 0.020 | 0.66 ± 0.020 | 0.65 ± 0.022 | 0.58 ± 0.019 |
| Figure 5B – 5F | *tenm4*$^{-/-}$ - Contrast = 0.05 | 0.04 ± 0.007 | 0.03 ± 0.010 | 0.04 ± 0.010 | 0.05 ± 0.009 | 0.03 ± 0.007 |
| Figure 5B – 5F | *tenm4*$^{-/-}$ - Contrast = 0.1 | 0.13 ± 0.012 | 0.12 ± 0.016 | 0.12 ± 0.015 | 0.16 ± 0.011 | 0.10 ± 0.015 |
| Figure 5B – 5F | *tenm4*$^{-/-}$ - Contrast = 0.25 | 0.38 ± 0.022 | 0.37 ± 0.022 | 0.38 ± 0.029 | 0.43 ± 0.023 | 0.32 ± 0.027 |
| Figure 5B – 5F | *tenm4*$^{-/-}$ - Contrast = 0.5 | 0.60 ± 0.028 | 0.57 ± 0.029 | 0.61 ± 0.036 | 0.62 ± 0.033 | 0.57 ± 0.027 |
| Figure 5B – 5F | *tenm4*$^{-/-}$ - Contrast = 1 | 0.67 ± 0.040 | 0.64 ± 0.051 | 0.70 ± 0.039 | 0.75 ± 0.046 | 0.60 ± 0.038 |
| Figure 5H | *tenm4*$^{+/+}$ - Bout Speed (1.3 – 20.6) | 1.89 ± 0.061 | 2.00 ± 0.045 | 2.24 ± 0.054 | 2.18 ± 0.047 | 2.04 ± 0.045 |
| Figure 5H | *tenm4*$^{+/-}$ - Bout Speed (1.3 – 20.6) | 1.93 ± 0.039 | 1.96 ± 0.039 | 2.19 ± 0.041 | 2.09 ± 0.038 | 2.05 ± 0.045 |
| Figure 5H | *tenm4*$^{-/-}$ - Bout Speed (1.3 – 20.6) | 1.85 ± 0.048 | 1.94 ± 0.043 | 2.18 ± 0.042 | 2.03 ± 0.040 | 1.96 ± 0.050 |
| Figure 5H | *tenm4*$^{+/+}$ - Bout Duration (1.3 – 20.6) | 136.18 ± 2.463 | 143.76 ± 2.231 | 160.04 ± 3.070 | 159.00 ± 3.109 | 151.61 ± 2.661 |

| | | | | | | |
|---|---|---|---|---|---|---|
| Figure 5H | *tenm4*$^{+/-}$ - Bout Duration (1.3 – 20.6) | 139.07 ± 1.721 | 143.65 ± 1.779 | 159.59 ± 2.052 | 153.14 ± 1.796 | 149.65 ± 2.117 |
| Figure 5H | *tenm4*$^{-/-}$ - Bout Duration (1.3 – 20.6) | 140.38 ± 2.171 | 144.74 ± 2.263 | 159.42 ± 2.783 | 154.12 ± 2.906 | 150.86 ± 2.738 |
| Figure 5H | *tenm4*$^{+/+}$ - Bout Distance (1.3 – 20.6) | 2.72 ± 0.125 | 3.02 ± 0.093 | 3.71 ± 0.123 | 3.64 ± 0.120 | 3.29 ± 0.102 |
| Figure 5H | *tenm4*$^{+/-}$ - Bout Distance (1.3 – 20.6) | 2.85 ± 0.081 | 2.99 ± 0.084 | 3.63 ± 0.091 | 3.36 ± 0.083 | 3.23 ± 0.099 |
| Figure 5H | *tenm4*$^{-/-}$ - Bout Distance (1.3 – 20.6) | 2.70 ± 0.092 | 2.94 ± 0.089 | 3.59 ± 0.083 | 3.26 ± 0.098 | 3.14 ± 0.116 |
| Figure 5H | *tenm4*$^{+/+}$ - Bout number (1.3 – 20.6) | 29.17 ± 0.803 | 31.37 ± 0.769 | 31.90 ± 0.743 | 29.84 ± 0.758 | 28.90 ± 0.813 |
| Figure 5H | *tenm4*$^{+/-}$ - Bout number (1.3 – 20.6) | 28.40 ± 0.611 | 30.68 ± 0.540 | 30.92 ± 0.544 | 29.37 ± 0.585 | 28.50 ± 0.526 |
| Figure 5H | *tenm4*$^{-/-}$ - Bout number (1.3 – 20.6) | 29.97 ± 0.892 | 32.35 ± 0.857 | 32.70 ± 0.948 | 30.98 ± 1.021 | 29.77 ± 0.966 |
| Figure 5H | *tenm4*$^{+/+}$ - Interbout Interval (1.3 – 20.6) | 0.60 ± 0.029 | 0.52 ± 0.022 | 0.49 ± 0.019 | 0.54 ± 0.023 | 0.58 ± 0.027 |
| Figure 5H | *tenm4*$^{+/-}$ - Interbout Interval (1.3 – 20.6) | 0.62 ± 0.023 | 0.54 ± 0.014 | 0.52 ± 0.014 | 0.56 ± 0.015 | 0.59 ± 0.018 |
| Figure 5H | *tenm4*$^{-/-}$ - Interbout Interval (1.3 – 20.6) | 0.54 ± 0.021 | 0.50 ± 0.018 | 0.46 ± 0.013 | 0.52 ± 0.017 | 0.55 ± 0.018 |
| Figure 5H | *tenm4*$^{+/+}$ - Correct Turns (1.3 – 20.6) | 55.21% ± 1.920% | 71.82% ± 1.735% | 79.33% ± 2.472% | 78.85% ± 2.523% | 75.63% ± 2.391% |
| Figure 5H | *tenm4*$^{+/-}$ - Correct Turns (1.3 – 20.6) | 53.61% ± 1.508% | 65.41% ± 1.966% | 76.43% ± 1.840% | 72.50% ± 2.054% | 70.71% ± 2.046% |
| Figure 5H | *tenm4*$^{-/-}$ - Correct Turns (1.3 – 20.6) | 52.81% ± 1.897% | 69.02% ± 2.863% | 76.34% ± 3.296% | 75.91% ± 3.495% | 75.18% ± 3.024% |
| Figure 5I | *tenm4*$^{+/+}$ - Bout Speed (1.3 – 20.6) | 1.87 ± 0.057 | 2.03 ± 0.057 | 2.27 ± 0.068 | 2.13 ± 0.061 | 2.05 ± 0.061 |
| Figure 5I | *tenm4*$^{+/-}$ - Bout Speed (1.3 – 20.6) | 1.94 ± 0.040 | 2.01 ± 0.040 | 2.25 ± 0.045 | 2.11 ± 0.039 | 1.96 ± 0.040 |
| Figure 5I | *tenm4*$^{-/-}$ - Bout Speed (1.3 – 20.6) | 1.93 ± 0.052 | 2.03 ± 0.052 | 2.21 ± 0.050 | 2.03 ± 0.051 | 1.91 ± 0.045 |
| Figure 5I | *tenm4*$^{+/+}$ - Bout Duration (1.3 – 20.6) | 136.53 ± 2.374 | 145.79 ± 2.472 | 161.24 ± 3.083 | 155.66 ± 3.118 | 152.38 ± 3.105 |
| Figure 5I | *tenm4*$^{+/-}$ - Bout Duration (1.3 – 20.6) | 140.66 ± 1.951 | 145.51 ± 1.821 | 162.29 ± 2.142 | 154.72 ± 1.790 | 150.22 ± 2.268 |
| Figure 5I | *tenm4*$^{-/-}$ - Bout Duration (1.3 – 20.6) | 143.22 ± 2.775 | 150.78 ± 3.136 | 161.64 ± 3.162 | 153.26 ± 3.120 | 146.68 ± 2.824 |
| Figure 5I | *tenm4*$^{+/+}$ - Bout Distance (1.3 – 20.6) | 2.73 ± 0.124 | 3.11 ± 0.117 | 3.87 ± 0.160 | 3.48 ± 0.139 | 3.31 ± 0.142 |
| Figure 5I | *tenm4*$^{+/-}$ - Bout Distance (1.3 – 20.6) | 2.89 ± 0.092 | 3.04 ± 0.079 | 3.82 ± 0.104 | 3.42 ± 0.081 | 3.08 ± 0.089 |
| Figure 5I | *tenm4*$^{-/-}$ - Bout Distance (1.3 – 20.6) | 2.92 ± 0.120 | 3.22 ± 0.127 | 3.76 ± 0.129 | 3.27 ± 0.116 | 2.97 ± 0.094 |
| Figure 5I | *tenm4*$^{+/+}$ - Bout number (1.3 – 20.6) | 29.59 ± 0.796 | 30.74 ± 0.762 | 30.82 ± 0.829 | 29.05 ± 0.802 | 27.93 ± 0.928 |
| Figure 5I | *tenm4*$^{+/-}$ - Bout number (1.3 – 20.6) | 28.16 ± 0.566 | 30.03 ± 0.501 | 30.99 ± 0.563 | 28.74 ± 0.563 | 27.15 ± 0.623 |

| | | | | | | |
|---|---|---|---|---|---|---|
| Figure 5I | *tenm4*$^{-/-}$ - Bout number (1.3 – 20.6) | 31.03 ± 0.671 | 32.29 ± 0.676 | 32.75 ± 0.829 | 31.14 ± 0.903 | 29.35 ± 0.851 |
| Figure 5I | *tenm4*$^{+/+}$ - Interbout Interval (1.3 – 20.6) | 0.58 ± 0.029 | 0.54 ± 0.022 | 0.52 ± 0.023 | 0.57 ± 0.027 | 0.63 ± 0.036 |
| Figure 5I | *tenm4*$^{+/-}$ - Interbout Interval (1.3 – 20.6) | 0.62 ± 0.020 | 0.55 ± 0.013 | 0.51 ± 0.014 | 0.57 ± 0.016 | 0.66 ± 0.025 |
| Figure 5I | *tenm4*$^{-/-}$ - Interbout Interval (1.3 – 20.6) | 0.52 ± 0.019 | 0.48 ± 0.015 | 0.47 ± 0.019 | 0.53 ± 0.022 | 0.58 ± 0.022 |
| Figure 5I | *tenm4*$^{+/+}$ - Correct Turns (1.3 – 20.6) | 56.48% ± 1.884% | 74.91% ± 1.817% | 82.19% ± 2.281% | 79.50% ± 2.514% | 77.80% ± 2.788% |
| Figure 5I | *tenm4*$^{+/-}$ - Correct Turns (1.3 – 20.6) | 55.73% ± 1.628% | 64.19% ± 1.927% | 78.66% ± 1.817% | 73.99% ± 2.258% | 74.08% ± 2.035% |
| Figure 5I | *tenm4*$^{-/-}$ - Correct Turns (1.3 – 20.6) | 55.94% ± 1.908% | 69.50% ± 2.987% | 78.89% ± 3.493% | 76.23% ± 3.339% | 76.60% ± 3.198% |
| Figure 5K (vertical left) | *tenm4*$^{+/+}$ - Bout Speed (1.3 – 20.6) | 1.90 ± 0.446 | 2.05 ± 0.356 | 2.28 ± 0.457 | 2.29 ± 0.403 | 2.08 ± 0.415 |
| Figure 5K (vertical left) | *tenm4*$^{+/-}$ - Bout Speed (1.3 – 20.6) | 1.92 ± 0.352 | 1.93 ± 0.343 | 2.13 ± 0.375 | 2.09 ± 0.298 | 2.01 ± 0.383 |
| Figure 5K (vertical left) | *tenm4*$^{-/-}$ - Bout Speed (1.3 – 20.6) | 1.85 ± 0.309 | 1.95 ± 0.356 | 2.23 ± 0.321 | 2.06 ± 0.332 | 2.00 ± 0.433 |
| Figure 5K (vertical left) | *tenm4*$^{+/+}$ - Bout Duration (1.3 – 20.6) | 135.05 ± 19.558 | 145.79 ± 17.578 | 161.06 ± 20.395 | 161.30 ± 21.405 | 153.34 ± 18.048 |
| Figure 5K (vertical left) | *tenm4*$^{+/-}$ - Bout Duration (1.3 – 20.6) | 138.17 ± 18.467 | 141.12 ± 18.232 | 155.22 ± 20.316 | 150.98 ± 17.884 | 149.52 ± 23.570 |
| Figure 5K (vertical left) | *tenm4*$^{-/-}$ - Bout Duration (1.3 – 20.6) | 139.18 ± 18.183 | 145.68 ± 18.263 | 164.72 ± 20.765 | 154.23 ± 20.610 | 147.28 ± 18.053 |
| Figure 5K (vertical left) | *tenm4*$^{+/+}$ - Bout Distance (1.3 – 20.6) | 2.70 ± 0.953 | 3.15 ± 0.785 | 3.82 ± 1.035 | 3.87 ± 0.977 | 3.39 ± 0.942 |
| Figure 5K (vertical left) | *tenm4*$^{+/-}$ - Bout Distance (1.3 – 20.6) | 2.82 ± 0.767 | 2.87 ± 0.730 | 3.44 ± 0.831 | 3.34 ± 0.758 | 3.20 ± 1.081 |
| Figure 5K (vertical left) | *tenm4*$^{-/-}$ - Bout Distance (1.3 – 20.6) | 2.68 ± 0.607 | 2.95 ± 0.736 | 3.80 ± 0.784 | 3.33 ± 0.824 | 3.14 ± 0.915 |
| Figure 5K (vertical left) | *tenm4*$^{+/+}$ - Bout number (1.3 – 20.6) | 29.81 ± 0.944 | 31.74 ± 0.866 | 32.77 ± 0.916 | 30.08 ± 0.912 | 29.06 ± 0.983 |
| Figure 5K (vertical left) | *tenm4*$^{+/-}$ - Bout number (1.3 – 20.6) | 28.63 ± 0.703 | 31.13 ± 0.636 | 31.39 ± 0.660 | 29.85 ± 0.730 | 29.06 ± 0.712 |
| Figure 5K (vertical left) | *tenm4*$^{-/-}$ - Bout number (1.3 – 20.6) | 30.29 ± 1.049 | 32.50 ± 1.091 | 32.92 ± 1.032 | 31.21 ± 1.153 | 29.67 ± 1.076 |
| Figure 5K (vertical left) | *tenm4*$^{+/+}$ - Interbout Interval (1.3 – 20.6) | 0.61 ± 0.231 | 0.52 ± 0.151 | 0.48 ± 0.137 | 0.54 ± 0.153 | 0.57 ± 0.168 |
| Figure 5K (vertical left) | *tenm4*$^{+/-}$ - Interbout Interval (1.3 – 20.6) | 0.62 ± 0.228 | 0.53 ± 0.112 | 0.52 ± 0.121 | 0.55 ± 0.139 | 0.59 ± 0.186 |
| Figure 5K (vertical left) | *tenm4*$^{-/-}$ - Interbout Interval (1.3 – 20.6) | 0.54 ± 0.144 | 0.50 ± 0.155 | 0.46 ± 0.116 | 0.53 ± 0.166 | 0.57 ± 0.163 |
| Figure 5K (vertical left) | *tenm4*$^{+/+}$ - Correct Turns (1.3 – 20.6) | 56.03% ± 2.352% | 72.69% ± 2.774% | 81.73% ± 3.074% | 81.10% ± 2.877% | 76.73% ± 3.333% |
| Figure 5K (vertical left) | *tenm4*$^{+/-}$ - Correct Turns (1.3 – 20.6) | 50.53% ± 2.118% | 62.75% ± 2.652% | 74.97% ± 2.810% | 70.47% ± 3.274% | 69.71% ± 2.774% |
| Figure 5K (vertical left) | *tenm4*$^{-/-}$ - Correct Turns (1.3 – 20.6) | 51.88% ± 2.772% | 69.30% ± 3.672% | 76.50% ± 4.260% | 76.51% ± 4.292% | 75.98% ± 3.976% |

| | | | | | | |
|---|---|---|---|---|---|---|
| Figure 5K (vertical right) | *tenm4*$^{+/+}$ - Bout Speed (1.3 – 20.6) | 1.88 ± 0.360 | 1.95 ± 0.282 | 2.19 ± 0.360 | 2.08 ± 0.311 | 2.00 ± 0.343 |
| Figure 5K (vertical right | *tenm4*$^{+/-}$ - Bout Speed (1.3 – 20.6) | 1.93 ± 0.354 | 1.99 ± 0.402 | 2.25 ± 0.452 | 2.10 ± 0.405 | 2.08 ± 0.448 |
| Figure 5K (vertical right | *tenm4*$^{-/-}$ - Bout Speed (1.3 – 20.6) | 1.85 ± 0.303 | 1.93 ± 0.280 | 2.12 ± 0.311 | 2.01 ± 0.260 | 1.92 ± 0.272 |
| Figure 5K (vertical right | *tenm4*$^{+/+}$ - Bout Duration (1.3 – 20.6) | 137.31 ± 16.154 | 141.74 ± 12.872 | 159.01 ± 20.027 | 156.71 ± 20.717 | 149.89 ± 20.158 |
| Figure 5K (vertical right | *tenm4*$^{+/-}$ - Bout Duration (1.3 – 20.6) | 139.98 ± 18.103 | 146.19 ± 21.055 | 163.97 ± 22.920 | 155.31 ± 21.022 | 149.78 ± 20.690 |
| Figure 5K (vertical right | *tenm4*$^{-/-}$ - Bout Duration (1.3 – 20.6) | 141.58 ± 17.022 | 143.81 ± 15.577 | 154.12 ± 17.911 | 154.01 ± 17.357 | 154.43 ± 20.556 |
| Figure 5K (vertical right | *tenm4*$^{+/+}$ - Bout Distance (1.3 – 20.6) | 2.73 ± 0.768 | 2.89 ± 0.532 | 3.59 ± 0.801 | 3.42 ± 0.782 | 3.19 ± 0.797 |
| Figure 5K (vertical right | *tenm4*$^{+/-}$ - Bout Distance (1.3 – 20.6) | 2.87 ± 0.781 | 3.11 ± 0.916 | 3.83 ± 1.082 | 3.38 ± 0.902 | 3.25 ± 0.930 |
| Figure 5K (vertical right | *tenm4*$^{-/-}$ - Bout Distance (1.3 – 20.6) | 2.73 ± 0.658 | 2.92 ± 0.634 | 3.39 ± 0.610 | 3.21 ± 0.555 | 3.15 ± 0.738 |
| Figure 5K (vertical right | *tenm4*$^{+/+}$ - Bout number (1.3 – 20.6) | 28.52 ± 0.961 | 31.00 ± 0.954 | 31.03 ± 1.006 | 29.60 ± 0.942 | 28.74 ± 1.043 |
| Figure 5K (vertical right | *tenm4*$^{+/-}$ - Bout number (1.3 – 20.6) | 28.18 ± 0.742 | 30.23 ± 0.639 | 30.44 ± 0.705 | 28.88 ± 0.750 | 27.93 ± 0.700 |
| Figure 5K (vertical right | *tenm4*$^{-/-}$ - Bout number (1.3 – 20.6) | 29.64 ± 1.022 | 32.19 ± 0.881 | 32.47 ± 1.161 | 30.76 ± 1.234 | 29.88 ± 1.132 |
| Figure 5K (vertical right | *tenm4*$^{+/+}$ - Interbout Interval (1.3 – 20.6) | 0.59 ± 0.177 | 0.51 ± 0.149 | 0.51 ± 0.140 | 0.55 ± 0.163 | 0.59 ± 0.202 |
| Figure 5K (vertical right | *tenm4*$^{+/-}$ - Interbout Interval (1.3 – 20.6) | 0.61 ± 0.186 | 0.54 ± 0.139 | 0.52 ± 0.136 | 0.58 ± 0.160 | 0.59 ± 0.171 |
| Figure 5K (vertical right | *tenm4*$^{-/-}$ - Interbout Interval (1.3 – 20.6) | 0.54 ± 0.144 | 0.49 ± 0.101 | 0.47 ± 0.110 | 0.50 ± 0.129 | 0.53 ± 0.140 |
| Figure 5K (vertical right | *tenm4*$^{+/+}$ - Correct Turns (1.3 – 20.6) | 54.38% ± 2.683% | 70.94% ± 2.721% | 76.94% ± 3.921% | 76.59% ± 3.724% | 74.52% ± 3.104% |
| Figure 5K (vertical right | *tenm4*$^{+/-}$ - Correct Turns (1.3 – 20.6) | 56.69% ± 2.271% | 68.79% ± 2.548% | 77.88% ± 2.947% | 74.52% ± 3.094% | 71.71% ± 2.765% |
| Figure 5K (vertical right | *tenm4*$^{-/-}$ - Correct Turns (1.3 – 20.6) | 53.75% ± 2.520% | 68.74% ± 3.790% | 76.19% ± 4.225% | 75.32% ± 4.523% | 74.38% ± 3.924% |
| Figure 5L (vertical left) | *tenm4*$^{+/+}$ - Bout Speed (1.3 – 20.6) | 1.86 ± 0.064 | 2.05 ± 0.061 | 2.31 ± 0.087 | 2.16 ± 0.081 | 2.14 ± 0.077 |
| Figure 5L (vertical left) | *tenm4*$^{+/-}$ - Bout Speed (1.3 – 20.6) | 1.89 ± 0.051 | 1.94 ± 0.056 | 2.22 ± 0.064 | 2.10 ± 0.055 | 1.91 ± 0.046 |
| Figure 5L (vertical left) | *tenm4*$^{-/-}$ - Bout Speed (1.3 – 20.6) | 1.91 ± 0.061 | 2.01 ± 0.079 | 2.17 ± 0.072 | 1.97 ± 0.065 | 1.90 ± 0.066 |
| Figure 5L (vertical left) | *tenm4*$^{+/+}$ - Bout Duration (1.3 – 20.6) | 136.20 ± 2.730 | 145.07 ± 2.813 | 161.22 ± 3.649 | 156.46 ± 4.295 | 153.82 ± 4.117 |
| Figure 5L (vertical left) | *tenm4*$^{+/-}$ - Bout Duration (1.3 – 20.6) | 140.64 ± 2.258 | 144.92 ± 2.416 | 159.76 ± 2.547 | 152.62 ± 2.238 | 147.49 ± 3.014 |
| Figure 5L (vertical left) | *tenm4*$^{-/-}$ - Bout Duration (1.3 – 20.6) | 144.00 ± 3.375 | 152.49 ± 4.471 | 163.40 ± 4.961 | 152.17 ± 4.763 | 145.94 ± 4.394 |
| Figure 5L (vertical left) | *tenm4*$^{+/+}$ - Bout Distance (1.3 – 20.6) | 2.68 ± 0.140 | 3.15 ± 0.135 | 3.94 ± 0.207 | 3.50 ± 0.164 | 3.49 ± 0.186 |

| | | | | | | |
|---|---|---|---|---|---|---|
| Figure 5L (vertical left) | *tenm4*$^{+/-}$ - Bout Distance (1.3 – 20.6) | 2.81 ± 0.115 | 2.91 ± 0.113 | 3.77 ± 0.151 | 3.37 ± 0.116 | 3.00 ± 0.113 |
| Figure 5L (vertical left) | *tenm4*$^{-/-}$ - Bout Distance (1.3 – 20.6) | 2.89 ± 0.140 | 3.21 ± 0.201 | 3.74 ± 0.212 | 3.15 ± 0.182 | 2.92 ± 0.160 |
| Figure 5L (vertical left) | *tenm4*$^{+/+}$ - Bout number (1.3 – 20.6) | 30.43 ± 0.921 | 31.16 ± 0.921 | 31.60 ± 1.006 | 30.16 ± 0.921 | 28.66 ± 1.016 |
| Figure 5L (vertical left) | *tenm4*$^{+/-}$ - Bout number (1.3 – 20.6) | 28.47 ± 0.709 | 29.97 ± 0.662 | 31.10 ± 0.711 | 29.26 ± 0.763 | 27.03 ± 0.744 |
| Figure 5L (vertical left) | *tenm4*$^{-/-}$ - Bout number (1.3 – 20.6) | 30.92 ± 1.090 | 32.48 ± 0.993 | 33.01 ± 1.181 | 30.76 ± 1.218 | 29.27 ± 1.145 |
| Figure 5L (vertical left) | *tenm4*$^{+/+}$ - Interbout Interval (1.3 – 20.6) | 0.55 ± 0.031 | 0.52 ± 0.025 | 0.50 ± 0.026 | 0.54 ± 0.028 | 0.59 ± 0.037 |
| Figure 5L (vertical left) | *tenm4*$^{+/-}$ - Interbout Interval (1.3 – 20.6) | 0.61 ± 0.022 | 0.55 ± 0.018 | 0.50 ± 0.018 | 0.56 ± 0.019 | 0.65 ± 0.029 |
| Figure 5L (vertical left) | *tenm4*$^{-/-}$ - Interbout Interval (1.3 – 20.6) | 0.53 ± 0.031 | 0.48 ± 0.020 | 0.45 ± 0.026 | 0.53 ± 0.030 | 0.57 ± 0.029 |
| Figure 5L (vertical left) | *tenm4*$^{+/+}$ - Correct Turns (1.3 – 20.6) | 53.95% ± 2.925% | 74.70% ± 2.496% | 83.19% ± 2.953% | 76.79% ± 3.603% | 75.40% ± 3.494% |
| Figure 5L (vertical left) | *tenm4*$^{+/-}$ - Correct Turns (1.3 – 20.6) | 53.20% ± 2.291% | 63.09% ± 2.387% | 78.66% ± 2.669% | 72.53% ± 3.084% | 72.66% ± 3.137% |
| Figure 5L (vertical left) | *tenm4*$^{-/-}$ - Correct Turns (1.3 – 20.6) | 55.92% ± 3.217% | 70.04% ± 3.534% | 77.58% ± 4.674% | 77.50% ± 4.381% | 75.66% ± 4.292% |
| Figure 5L (vertical right) | *tenm4*$^{+/+}$ - Bout Speed (1.3 – 20.6), | 1.88 ± 0.065 | 2.01 ± 0.068 | 2.22 ± 0.080 | 2.10 ± 0.069 | 1.95 ± 0.076 |
| Figure 5L (vertical right) | *tenm4*$^{+/-}$ - Bout Speed (1.3 – 20.6) | 1.99 ± 0.058 | 2.08 ± 0.053 | 2.28 ± 0.061 | 2.12 ± 0.057 | 2.00 ± 0.070 |
| Figure 5L (vertical right) | *tenm4*$^{-/-}$ - Bout Speed (1.3 – 20.6) | 1.96 ± 0.059 | 2.06 ± 0.065 | 2.25 ± 0.074 | 2.09 ± 0.063 | 1.92 ± 0.061 |
| Figure 5L (vertical right) | *tenm4*$^{+/+}$ - Bout Duration (1.3 – 20.6) | 136.86 ± 2.898 | 146.50 ± 3.022 | 161.25 ± 4.347 | 154.85 ± 3.588 | 150.95 ± 4.532 |
| Figure 5L (vertical right) | *tenm4*$^{+/-}$ - Bout Duration (1.3 – 20.6) | 140.68 ± 2.916 | 146.10 ± 2.628 | 164.82 ± 3.503 | 156.82 ± 3.296 | 152.96 ± 3.775 |
| Figure 5L (vertical right) | *tenm4*$^{-/-}$ - Bout Duration (1.3 – 20.6) | 142.45 ± 3.841 | 149.07 ± 3.716 | 159.88 ± 4.054 | 154.34 ± 4.042 | 147.42 ± 3.265 |
| Figure 5L (vertical right) | *tenm4*$^{+/+}$ - Bout Distance (1.3 – 20.6) | 2.77 ± 0.151 | 3.07 ± 0.152 | 3.79 ± 0.230 | 3.45 ± 0.176 | 3.14 ± 0.195 |
| Figure 5L (vertical right) | *tenm4*$^{+/-}$ - Bout Distance (1.3 – 20.6) | 2.96 ± 0.139 | 3.17 ± 0.114 | 3.86 ± 0.156 | 3.48 ± 0.141 | 3.15 ± 0.154 |
| Figure 5L (vertical right) | *tenm4*$^{-/-}$ - Bout Distance (1.3 – 20.6) | 2.95 ± 0.140 | 3.23 ± 0.147 | 3.78 ± 0.183 | 3.38 ± 0.157 | 3.01 ± 0.121 |
| Figure 5L (vertical right) | *tenm4*$^{+/+}$ - Bout number (1.3 – 20.6) | 28.74 ± 0.906 | 30.32 ± 0.890 | 30.04 ± 1.083 | 27.94 ± 1.007 | 27.20 ± 1.084 |
| Figure 5L (vertical right) | *tenm4*$^{+/-}$ - Bout number (1.3 – 20.6) | 27.86 ± 0.744 | 30.09 ± 0.623 | 30.87 ± 0.769 | 28.22 ± 0.781 | 27.27 ± 0.895 |
| Figure 5L (vertical right) | *tenm4*$^{-/-}$ - Bout number (1.3 – 20.6) | 31.13 ± 0.923 | 32.10 ± 0.843 | 32.50 ± 1.094 | 31.53 ± 1.144 | 29.42 ± 1.135 |
| Figure 5L (vertical right) | *tenm4*$^{+/+}$ - Interbout Interval (1.3 – 20.6) | 0.61 ± 0.034 | 0.55 ± 0.026 | 0.55 ± 0.034 | 0.60 ± 0.035 | 0.67 ± 0.041 |
| Figure 5L (vertical right) | *tenm4*$^{+/-}$ - Interbout Interval (1.3 – 20.6) | 0.63 ± 0.031 | 0.54 ± 0.019 | 0.51 ± 0.021 | 0.58 ± 0.025 | 0.67 ± 0.040 |

| | | | | | | |
|---|---|---|---|---|---|---|
| Figure 5L (vertical right) | *tenm4*$^{-/-}$ - Interbout Interval (1.3 – 20.6) | 0.51 ± 0.023 | 0.49 ± 0.019 | 0.49 ± 0.024 | 0.52 ± 0.030 | 0.59 ± 0.030 |
| Figure 5L (vertical right) | *tenm4*$^{+/+}$ - Correct Turns (1.3 – 20.6) | 59.00% ± 2.860% | 75.12% ± 2.475% | 81.19% ± 2.783% | 82.21% ± 2.638% | 80.20% ± 3.358% |
| Figure 5L (vertical right) | *tenm4*$^{+/-}$ - Correct Turns (1.3 – 20.6) | 58.40% ± 2.494% | 64.84% ± 2.679% | 78.02% ± 2.945% | 74.81% ± 3.249% | 75.56% ± 3.019% |
| Figure 5L (vertical right) | *tenm4*$^{-/-}$ - Correct Turns (1.3 – 20.6) | 55.97% ± 2.493% | 68.96% ± 3.755% | 80.21% ± 3.963% | 74.96% ± 4.067% | 77.54% ± 3.876% |
